# EPH Receptor Tyrosine Kinases Regulate Epithelial Morphogenesis and Phosphorylate the PAR-3 Scaffold Protein to Modulate Downstream Signaling Networks

**DOI:** 10.1101/2020.09.06.285270

**Authors:** Sara L. Banerjee, Noémie Lavoie, Kévin Jacquet, Frédéric Lessard, Ana Osornio-Hernandez, Josée N. Lavoie, Sabine Elowe, Jean-Philippe Lambert, Patrick Laprise, Nicolas Bisson

## Abstract

The EPH family is the largest among receptor tyrosine kinases (RTKs) in humans. In contrast to other RTKs, EPH receptors (EPHRs) cognate ligands, ephrins, are tethered to the cell surface. This results in EPHR-ephrin signaling being mainly involved in short-range cell-cell communication events that regulate cell adhesion, migration and tissue boundary formation. Although EPHRs functions have been broadly studied, the molecular mechanisms by which they control these processes are far from being understood. To address this, we sought to identify new effector proteins acting downstream of EPHRs and determine their role in EPHR-regulated functions. To unravel EPHR-associated signaling complexes under native conditions, we applied a mass spectrometry-based approach, namely BioID proximity labeling. We obtained a composite proximity network from EPHA4, -B2, -B3 and -B4 receptors that comprises 395 proteins, most of which were not previously linked to EPH signaling. A gene ontology and pathway term analysis of the most common candidates highlighted cell polarity as a novel function associated with EPHR activity. We found that EPHA1 and EPHB4 expression is restricted to the basal and lateral membrane domains in polarized Caco-2 3D spheroidal cell cultures. We further discovered that their depletion impairs the compartmentalized distribution of polarity proteins as well as overall spheroid morphogenesis. Moreover, we examined the contribution of a number of candidates, selected from EPHR proximity networks, via loss-of-function in an EPHR-dependent cell segregation assay. We found that depletion of the signaling scaffold PAR-3 blocks cell sorting. We also delineated a signalling complex involving the C-terminal SRC kinase (CSK), whose recruitment to PAR-3 complexes is dependent on EPHR signals. Our work sheds a new light on EPHR signaling networks and describes conceptually novel the mechanisms by which EPHRs signal at the membrane to contribute to the regulation of cellular phenotypes.

## INTRODUCTION

Cells communicate with each other to ensure proper development and function of multicellular organisms. Cell-cell communication is often mediated by transmembrane receptors that trigger signaling networks to dictate the response to extracellular cues (Hunter 2000, Scott and Pawson 2009, Nair et al. 2019). The EPH family of receptor tyrosine kinases (RTKs) is composed of fourteen members that interact with membrane-tethered ephrin ligands (Lemmon and Schlessinger 2010). The EPHR/ephrin-mediated cell-cell contact-dependent communication is essential for a variety of biological processes that require highly defined spatial guidance for cell positioning in many tissues of the developing and adult organism (Kania and Klein 2016). Extensively documented examples of the effect of EPHR activation following ligand binding comprise the regulation of cell migration, repulsion and cell-cell adhesion during axonal pathfinding and tissue boundary formation (Niethamer and Bush 2019). Most EPHRs are found overexpressed in a variety of human tumors, especially in more aggressive and lethal ones (Fleuren et al. 2016, Kou and Kandpal 2018). Although the correlation of EPH overexpression with poor disease outcome in cancer patients argues for these receptors acting as tumor promoting factors, accumulating evidence suggests that they can also function as tumor suppressors (Pasquale 2010, Buckens et al. 2020). While a number of studies have unveiled factors contributing to EPHR signaling complexity (as reviewed in (Lisabeth, Falivelli, and Pasquale 2013, Nikolov, Xu, and Himanen 2013, Kania and Klein 2016)), intracellular signaling networks that are essential for EPHR ability to control cell phenotype remain vastly unexplored. The list of EPHR effectors is largely incomplete, raising questions about how EPHRs execute their diverse biological functions. Collectively, this limits our understanding of EPHR signaling systems in both normal physiology and disease states (Pasquale 2010).

A global identification of EPHR-centric protein complexes may shed light on core EPHRs signaling networks, as well as allowing to decipher specific signaling outputs from selected receptors. This would significantly contribute to addressing how distinct EPHRs transmit signals to achieve diverse biological outcomes. Affinity purification coupled to mass spectrometry (AP-MS) has been a primary approach for the identification of macromolecular signaling complexes and analyses of protein interaction networks that occur in living cells (Gingras et al. 2007). However, AP-MS requires preserving protein interactions (PPIs) throughout the purification process, which generally rely on bait protein solubility (Gingras and Raught 2012). As a result, mapping interactions for plasma membrane resident proteins such as EPHRs has proven challenging (Gong et al. 2016, Huttlin et al. 2020). The development of proximity labeling approaches combined to MS, for example BioID (Roux et al. 2012), overcomes these limitations of conventional AP-MS (Gingras, Abe, and Raught 2019). BioID allows mapping proximal, transient or stable interactions while they are taking place in their native cellular environment. This strategy has already been applied for the efficient mapping of neighboring proteins from different subcellular compartments, including the plasma membrane (Jacquet et al. 2018, Lambert et al. 2019, Go et al. 2019).

Here, we used BioID to unravel a high-confidence EPHR proximity signaling network comprising 395 candidate proteins. Gene ontology analyses highlighted cell polarity as a putative novel function for EPH signaling. We described a compartmentalized distribution of EPHRs in polarized epithelial cells 3D cysts and revealed the requirement for EPHA1 and EPHB4 in morphogenesis, which is intimately linked to cell polarity. We further explored the contribution of a number of BioID-identified candidates in EPH-mediated cell sorting. We found that the polarity scaffold protein PAR-3 is a substrate for EPHA4, which is required for cell segregation, in a signaling complex that relies on the pTyr-dependent recruitment of CSK. Thus, our work sheds light on an unexplored function for EPHRs in cell polarity, as well as on a new EPHR-induced signaling complex containing PAR-3.

## RESULTS

### BioID proteomics delineate EPHR proximity signaling networks

To identify new putative downstream effector proteins for EPHRs, we selected four EPHRs responding to the ephrin-B type ligands, namely EPHA4, -B2, -B3 and -B4. We fused BirA* (Roux et al. 2012), a promiscuous mutant biotin ligase that catalyses covalent biotin binding to Lys residues on adjacent proteins, to each receptor C-terminus. We chose the HEK293 cell line, as it has previously proven useful for the study of EPH-controlled biological processes (Jørgensen et al. 2009) and displays endogenous expression of most human EPHRs (Schweppe et al. 2018, Huttlin et al. 2020). Using the Flp-In T-Rex system, we generated inducible HEK293 T-Rex stable cell lines expressing EPHR-BirA*-FLAG baits or a YFP-BirA*-FLAG control. We verified the inducible expression and the relative protein levels of the BirA* chimeric fusions following tetracycline induction (Figure 1A). We also validated the compartmentalized enrichment of all EPHR fusions at the plasma membrane (Figure 1B). To determine whether the chimeras maintained functional signaling (EPHR) and biotin ligase (BirA*) activities, we performed two sets of experiments. First, we tested whether EPHRs were activated in an ephrin ligand-dependent manner. EPHR-BirA*-FLAG cells were mixed with ephrinB1 (EFNB1)-(for EPHB2, -B3) or EFNB2-expressing cells (for EPHA4, -B4) for 15 or 30 minutes, and activation was assessed by receptor autophosphorylation. All four chimeras displayed a notable increase in Tyr phosphorylation as soon as 15 minutes following mixing with ligand-expressing cells (Figure 1C). Second, we assessed the biotin ligase activity of BirA*. Addition of exogenous biotin to the cell culture medium led to a strong increase in biotinylation of endogenous proteins compared to controls, as revealed by Streptavidin Western blotting (Figure 1D). Together, these results indicate that EPHR-BirA*-FLAG fusions maintain the receptors’ plasma membrane compartmentalization and their signaling and biotinylation activities.

**Figure 1.**
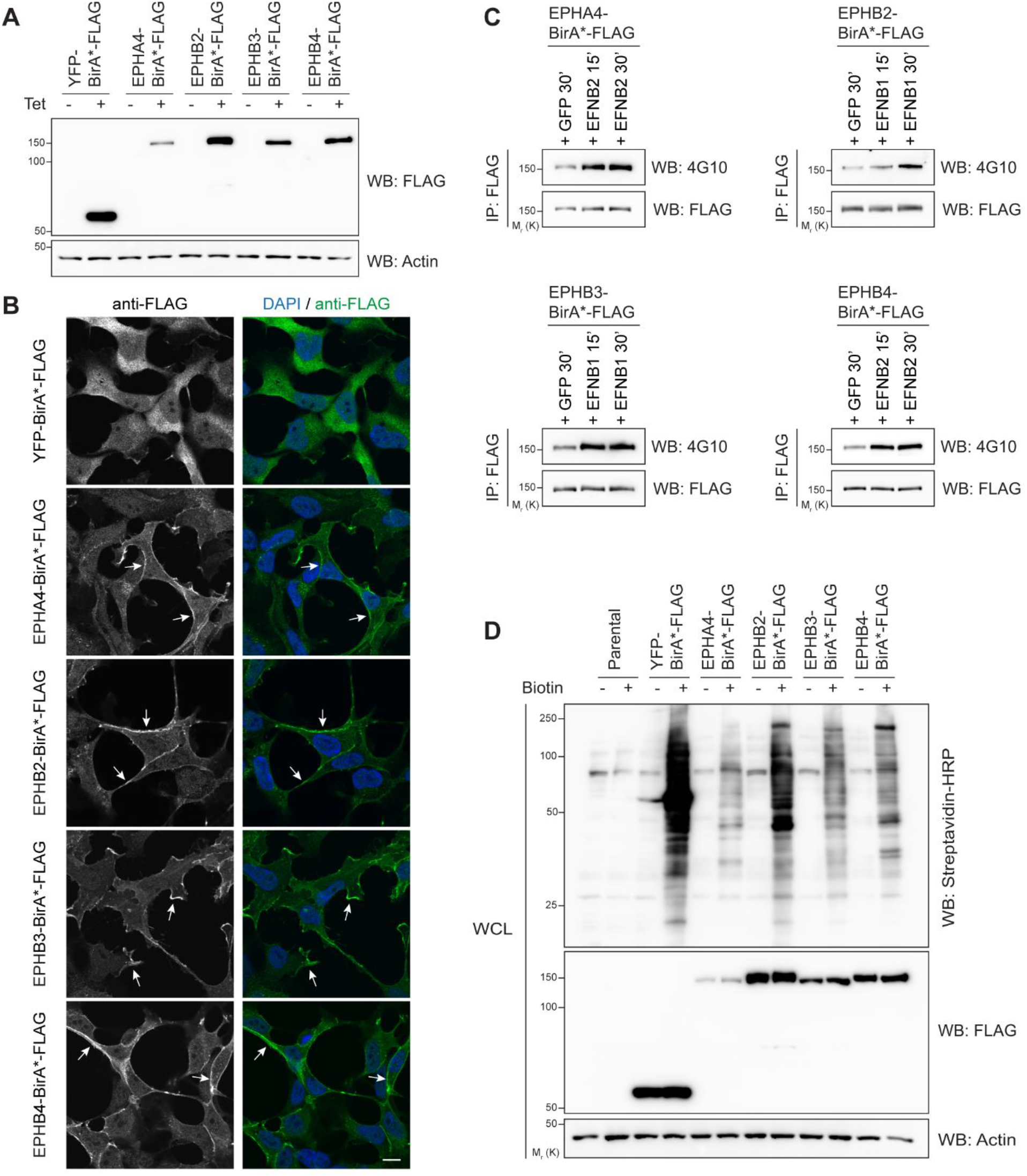
HEK293 stable cells express functional EPHR-BirA*-FLAG fusion proteins. (A) Western blot analysis of the inducible expression of EPHR-BirA*-FLAG fusions in HEK293 T-Rex cells. (B) Confocal images displaying the subcellular compartmentalization of BirA* chimeras (scale bar: 10 μm). The EPHR-BirA*-FLAG plasma membrane enrichment is indicated with arrows. (C) Western blot analysis displaying Tyr (auto)phosphorylation of EPHR-BirA*-FLAG fusions following 15 or 30 minutes mixing with HEK293 cells expressing cognate ephrin ligands (EFNB1 or EFNB2). (D) Analysis of total endogenous protein biotinylation following addition of biotin for 24 hours to the cell culture medium, as revealed by Streptavidin-HRP Western blotting.

To dissect EPHR proximity signaling networks, we carried protein labeling with an excess of biotin for 24 hours. We then, performed streptavidin affinity-purification of biotinylated proteins from cell lysates, followed by their identification via LC-MS/MS. Three independent experiments were completed for each EPHR-BirA*-FLAG fusion, as well as the YFP-BirA*-FLAG control. We eliminated background contaminants and non-specific interactions via the SAINTexpress (significance analysis of interactome) algorithm (Teo et al. 2014), by scoring against the YFP-BirA*-FLAG control and using a stringent SAINT Bayesian false-discovery rate (BFDR) ≤1% as a threshold (Figure S1 and Table S1). This statistical analysis revealed 307, 125, 236 and 193 high-confidence proximity partners for EPHA4, -B2, -B3 and -B4, respectively, for a total of 395 non-redundant interaction partners (Figures 2A and S2). Our BioID experiments identified a number of well-known EPHR signaling effectors (Figures S1, S2). This group includes kinases YES1 (Zisch et al. 1998), ABL2 (Yu et al. 2001) and subunit PIK3R1 (Pandey et al. 1994, Fang et al. 2008), adaptor proteins NCK2 (Bisson et al. 2007, Fawcett et al. 2007, Tanaka et al. 2003) and SHB (Wagner et al. 2020), as well as RHO family GTPases guanine nucleotide exchange factors (GEFs) ITSN1 (Irie and Yamaguchi 2002) and TIAM1 (Tolias et al. 2007). We also identified EFNB1 and EFNB2 ligands (Noberini, Rubio de la Torre, and Pasquale 2012). This confirmed the validity of our proximity labeling approach. The identification of the previously characterized EPHR PDZ motif-dependent partner AFDN (Hock et al. 1998, Beaudoin et al. 2012) also implied that the C-terminal BirA* fusion did not impair motif accessibility to cognate PDZ domains.

**Figure 2.**
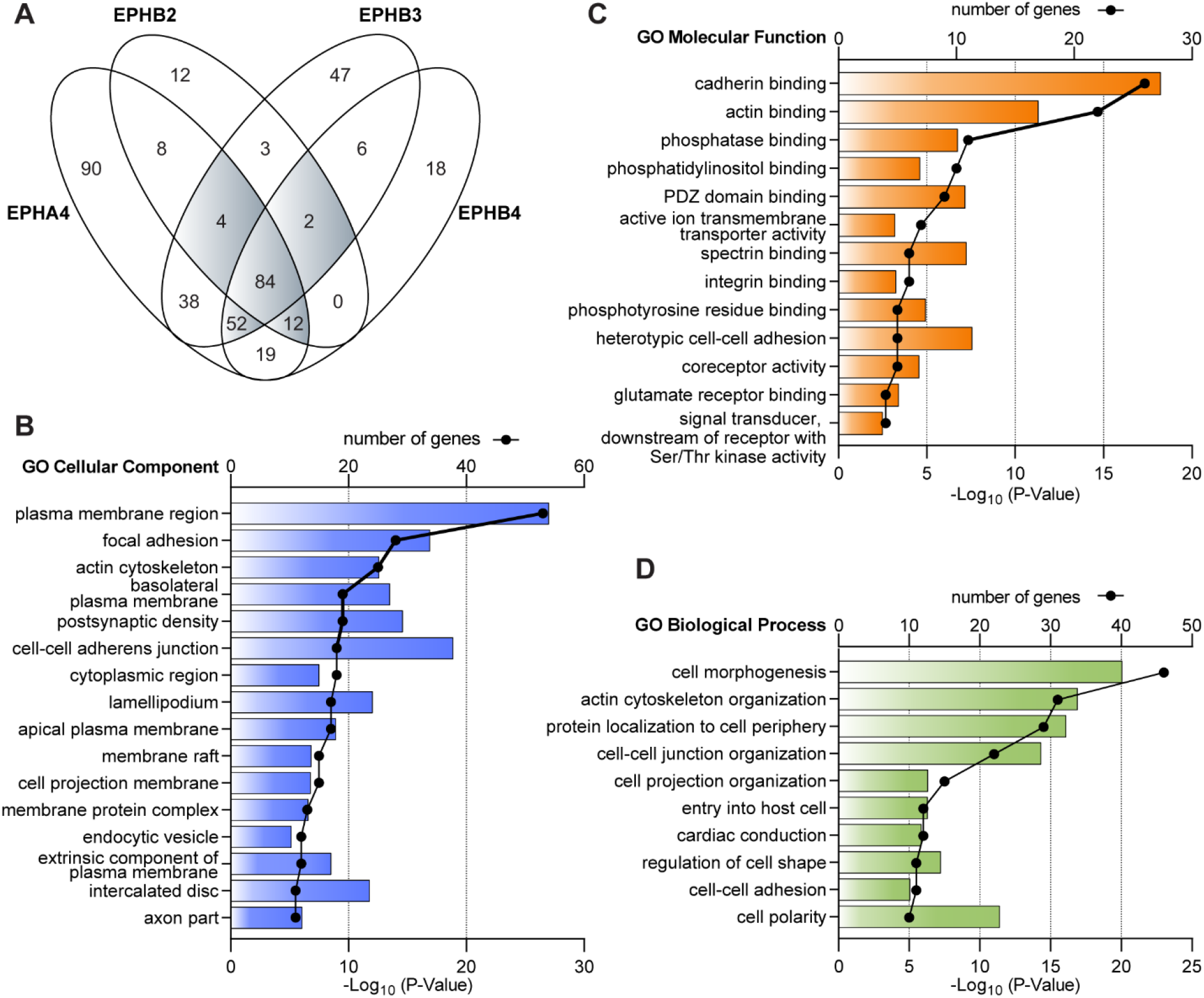
BioID proximity labeling proteomics reveal both known and unexplored functions for EPHRs. (A) Venn diagram representing the number of *bona fide* components of EPHA4, EPHB2, EPHB3 and EPHB4 proximity networks following the removal of background contaminants and non-specific interactions via SAINTexpress using a YFP-BirA*-FLAG control (BFDR <1%). Proteins found with at least three EPHRs are highlighted in grey. Individual identifications are provided in Figure S2. (B-D) Bar graph representation of gene ontology (GO) analysis via ClueGo of cellular components (panel B), molecular functions (panel C) and biological processes (panel D) clusters linked with at least 10 candidates from the group shaded in grey in panel A.

While nearly half (n=167, 42%) of the 395 proteins that we identified were labeled as being specific to a given EPHR among the four that we surveyed, a significant number of candidates were deemed common to at least three (n=154, 39%) (Figures 2A and S2). This shed light on signaling complexes that are potentially required for EPHRs to execute core biological functions. Using this group as a reference, we performed gene ontology (GO) analysis to associate specialized molecular functions and biological processes with components of the EPHR proximity network (Bindea et al. 2009). Our analysis revealed a strong enrichment for plasma membrane-targeted cellular components, as well as cadherin and actin binding properties (Figures 2B and 2C). Additionally, it highlighted the regulation of cell morphogenesis and actin cytoskeleton organization as the main processes, with protein localization to cell periphery, cardiac conduction and entry into host cells also emerging among the top groups (Figure 2D). Accordingly, previous reports have implicated EPH signaling in these biological processes (Barquilla and Pasquale 2015, Kania and Klein 2016), further supporting the validity of our BioID data. We also found 10 candidates associated with the regulation of cell polarity (p<1e10), a process not previously linked to EPH signaling (Figure 2D). Collectively, these results suggest that EPHRs may participate in establishing cell polarity, or use polarity proteins as novel effectors to control previously established EPHR-driven functions.

### EPHA1 or EPHB4 depletion impairs the establishment of cell morphogenesis and polarity in a 3D spheroid model

To evaluate the relative contribution of EPHRs to tissue morphogenesis and polarity, we opted for Caco-2 human colorectal adenocarcinoma cells. Following seeding in Matrigel, these cells form spheroidal cysts consisting of an epithelial monolayer where the apical surface faces a single hollow lumen (Jaffe et al. 2008). We surveyed the Human Protein Atlas for EPHR gene expression in Caco-2 cells and found *EPHA1, -A2, -A4* and *-B4* to be the most abundantly expressed (Figure S3A) (Uhlén et al. 2015). Their expression at the protein level was confirmed by MS-based analysis of Caco-2 cells proteome (Zecha et al. 2020). Using Western blotting, we assessed whether these four EPHRs were expressed in confluent and post-confluent Caco-2 cells, as they begin to differentiate and to establish apical-basal polarity once they reach confluence (Jaffe et al. 2008). We detected all four receptors in sub-confluent cells, but only EPHA1 and EPHB4 expression was sustained at confluency and post-confluency in monolayers (Figure S3B), suggesting a potential role in the establishment of cell polarity. For this reason, we pursued our experiments in Caco-2 cells with these 2 receptors. In particular, the EPHB4 proximity network revealed a putative association with >10 polarity proteins, including PAR-3, PAR-6B and PKCι (Figure S2). In 3D spheroidal cysts, EPHA1 and EPHB4 both localized at the lateral and basal membranes (Figure 3A). This was evidenced by signals that were restricted basal to the ZO-1 (TJP1) tight junction marker and that did not coincide with apical Ezrin (EZR) staining. The compartmentalized distribution of both EPHA1 and EPHB4 suggested that these receptors are part of the polarity networks that control tissue morphogenesis.

**Figure 3.**
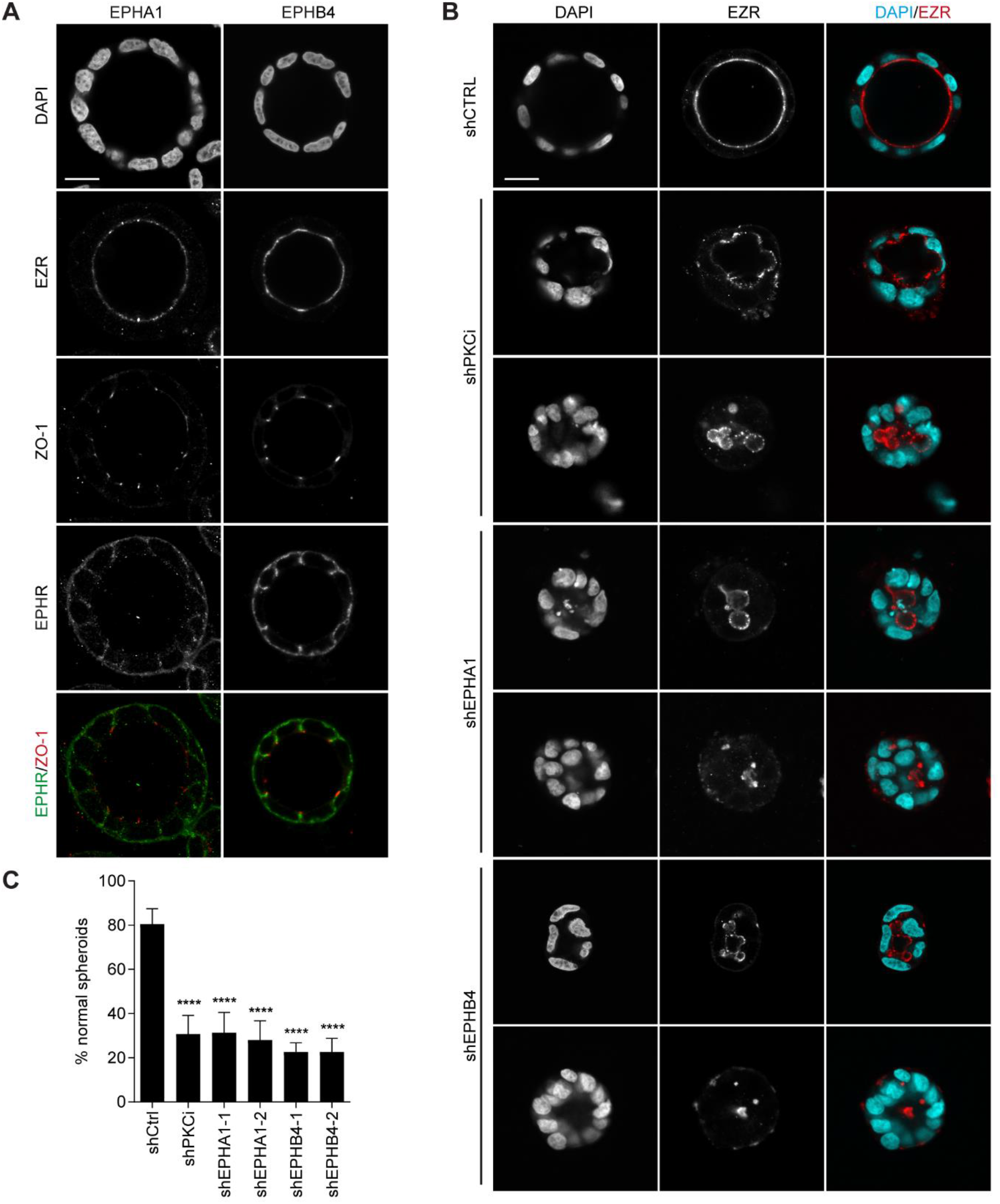
EPHA1 and EPHB4 display a compartmentalized localization in Caco-2 spheroids and are required for adequate morphogenesis. (A) Confocal images displaying a representative example of EPHA1 and EPHB4 expression in 3D Caco-2 spheroid cysts. Single cells were seeded on Matrigel and grown for 6 days. Expression of apical (EZR) and tight junction (ZO-1) markers is shown, along with nuclear DAPI staining (scale bar: 15 µm). (B) Representative images of Caco-2 spheroids morphology following depletion of PKCι, EPHA1 or EPHB4. Two independent shRNAs were used against EPHRs. EZR apical marker highlights the lumen border. Cells were grown as in (A) (scale bar: 15 µm). (C) Quantification of the percentage of Caco-2 spheroids displaying a wild-type morphology (i.e. single, round lumen judged by EZR staining, surrounded by a monolayer of cells) following depletion of PKCι, EPHA1 or EPHB4. The mean of three independent experimental replicates is shown. Error bars indicate SD (****p< 0.0001; unpaired t-test; n=450 for shCTRL and shPKCι, and n=150 for each shEPH).

To test whether EPHA1 and EPHB4 are required for the morphogenesis of polarized spheroids, we performed shRNA-mediated EPHR depletion in Caco-2 cells prior to seeding in Matrigel (Figure 3B). We achieved a >85% depletion using two distinct shRNA sequences for each EPHR tested (Figure S4). We found that EPHA1 or EPHB4 depletion led to the formation of disrupted spheroid structures, similar to those obtained following the depletion of the polarity protein PKCι (PRKCI) (Figure 3B). We observed that >70% randomly analyzed EPHA1- or EPHB4-depleted spheroids displayed aberrant apical EZR staining that highlighted the presence of multiple apical loci or a disrupted apical surface (n>150, p<0.0001; unpaired t-test) (Figure 3C). The penetrance of this aberrant phenotype was equivalent to that achieved with PKCι depletion (Figures 3C and S4). Overall, our results highlight the compartmentalized expression of EPHA1 and EPHB4 in polarized epithelial spheroidal cysts and support a role for these two tyrosine kinase receptors in tissue polarity and morphogenesis, as suggested by the identification of multiple polarity proteins in the EPHR proximity network.

### PAR-3 polarity protein is required for EPHR-dependent cell segregation

Our BioID proteomic experiments also revealed a number of putative effectors, including polarity proteins that may contribute to EPHR-driven cellular phenotypes (Figure S2). To evaluate whether these candidates are required for EPHR signaling, we examined the effect of their depletion using a well-established EFN ligand-dependent, EPHR-driven cell segregation assay that is reminiscent of tissue boundary formation *in vivo* (Figure 4A) (Wu et al. 2019). Co-culture of EPHR-expressing GFP-labeled HEK293 cells with cells expressing a cognate EFNB1 ligand led to the formation of large homogenous clusters of EPHR-expressing cells (Figure 4A-B), a consequence of EPH-ephrin contact restricting the two cell types from intermingling. Conversely, co-culture of the same EPH-expressing cells with a parental line resulted in cells mixing uniformly, a condition that we used as a negative control (Figure 4A-B). We selected 11 candidates that were identified in proximity networks of at least three of the four EPHRs used as baits (Figures 2A and S2), as well as 5 additional targets that we picked due to their homology to the candidates (PARD3B) or their reported association with EPHRs: GRB2 (Moeller et al. 2006, Stein, Cerretti, and Daniel 1996), NCK1 (Stein et al. 1998, Hu et al. 2009), NCK2 (Bisson et al. 2007, Fawcett et al. 2007, Tanaka et al. 2003) and PTPN11 (Miao et al. 2000, Miura et al. 2013). We individually depleted these 16 proteins by transfecting two distinct siRNAs specifically in the EPHR-expressing cells prior to co-culture, thus ensuring that any phenotype would be autonomous to the EPHR-expressing population. Using average cluster size as a quantitative measure to evaluate cell segregation, we found that 5/16 candidates significantly impaired the EPHR-dependent clustering (p<0.0001 for both siRNAs; unpaired t-test) (Figures 4C and S5). Depletion of previously characterized EPHR effector NCK2 led to a block of cell segregation, resulting in reduced cluster formation. Among the tested BioID candidates, depletion of the scaffold/polarity protein PAR-3 (PARD3) led to the strongest and most consistent reduction in cluster size relative to control, while no effect was measured when depleting another polarity protein, SCRIB (Figures. 4B and 4C). To further validate this observation, we depleted PAR-3 using a shRNA targeting a different sequence on *PARD3* mRNA and demonstrated that the PAR-3 loss-of-function blocked EPH-dependent cell sorting (p<0.0001; unpaired t-test), thus confirming PAR-3 requirement for the phenotype (Figures S6A-C). Our data suggest that PAR-3 is a novel effector of EPHR signaling.

**Figure 4.**
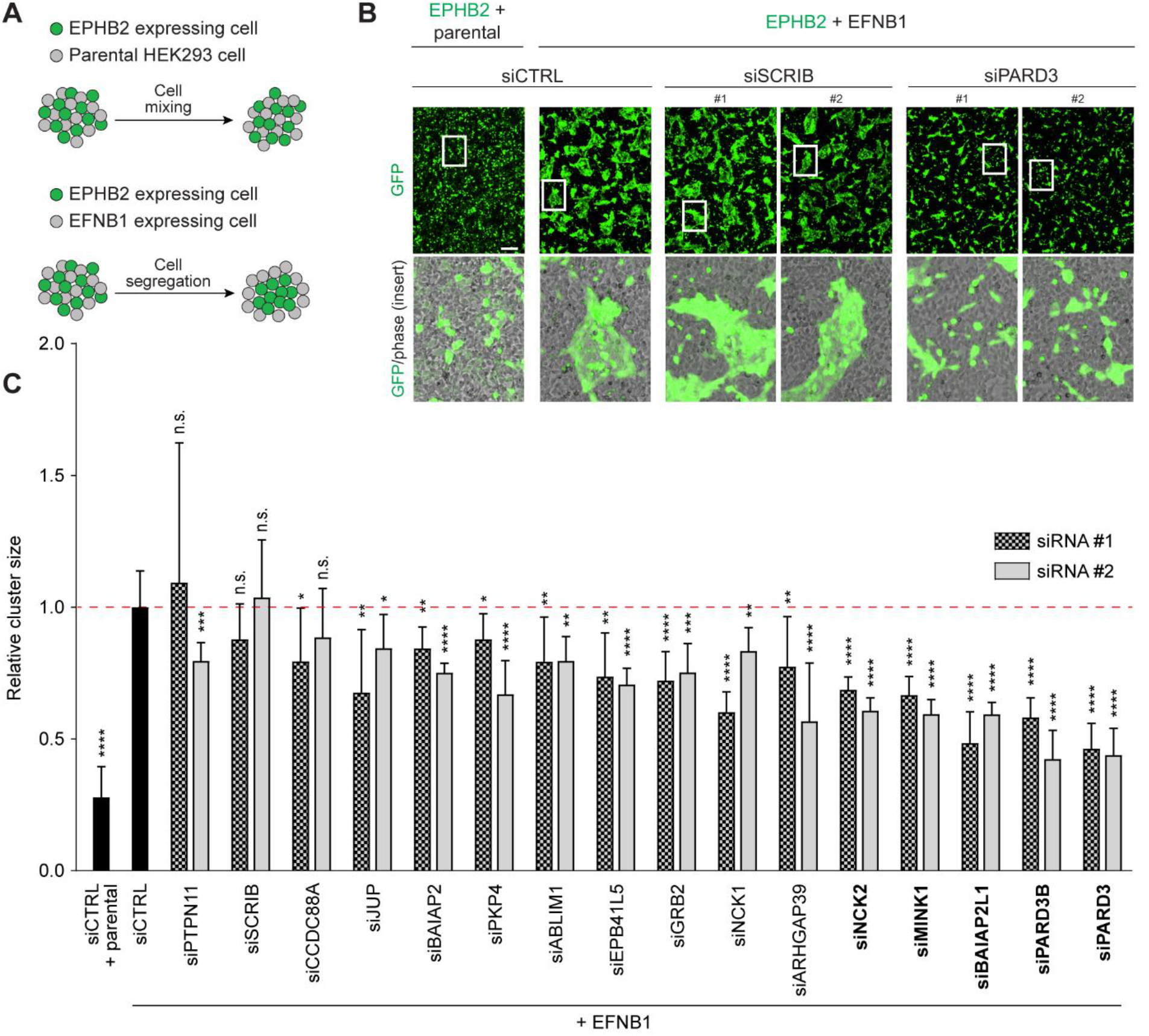
PAR-3 polarity protein is required for EPHR-dependent cell segregation. (A) Schematic of the EPHB2-/EFNB1-dependent co-culture cell segregation assay employed for functional examination of selected BioID candidate hits. (B) Representative images of cell co-cultures following siRNA depletion of SCRIB or PAR-3 (PARD3) in the EPHB2 expressing cells, visualized by GFP (scale bar: 250 µm). (C) Quantification of the average size of EPHB2 GFP+ cell clusters depleted with selected candidates with two independent siRNAs, following mixing with ephrinB1-expressing cells, relative to control (EPHB2 siCTRL cells mixed with ephrinB1 cells). The mean of two independent experimental replicates is shown. Error bars indicate SD (*p < 0.05; **p < 0.01; *** p < 0.001; **** p < 0.0001; unpaired t-test). Proteins for which depletion via both siRNAs tested resulted in the strongest and most consistent reduction in cluster size relative to control (p<0.0001; unpaired t-test) are in bold.

### EPHA4 associates with and phosphorylates PAR-3

To better define the relationship between EPHRs and PAR-3, we assessed whether the two proteins are present in the same signaling complexes, as BioID enables both direct and indirect (proximity dependent) protein-protein interactions to be identified. To explore this possibility, we transiently transfected GFP-labelled PAR-3 or a GFP control in our EPHR-BirA*-FLAG lines and performed anti-GFP affinity purifications followed by Western blotting. We found that EPHA4, -B2, -B3 and -B4 specifically co-purified with PAR-3 but not with our GFP control (Figure 5A). We also discovered that EPHR overexpression coincides with PAR-3 Tyr phosphorylation. We opted to focus on the PAR-3-EPHA4 pair as this receptor yielded the largest number of targets in the BioID experiments (Figure S2), and because catalytically active EPHA4 can be purified to homogeneity (Dionne et al. 2018). To validate the association with PAR-3 in absence of the BirA*-FLAG fusion, we co-transfected GFP-PAR-3 with wild-type (WT) EPHA4 into HEK293T cells and performed affinity purification followed by Western blot analysis. We confirmed the association between PAR-3 and EPHA4. The receptor catalytic activity was not required for the interaction, but was essential for PAR-3 Tyr phosphorylation, as demonstrated using a kinase inactive mutant (kinase dead (KD); K653A) (Figure 5B). These observations suggested that PAR-3 may be a substrate for EPHA4.

**Figure 5.**
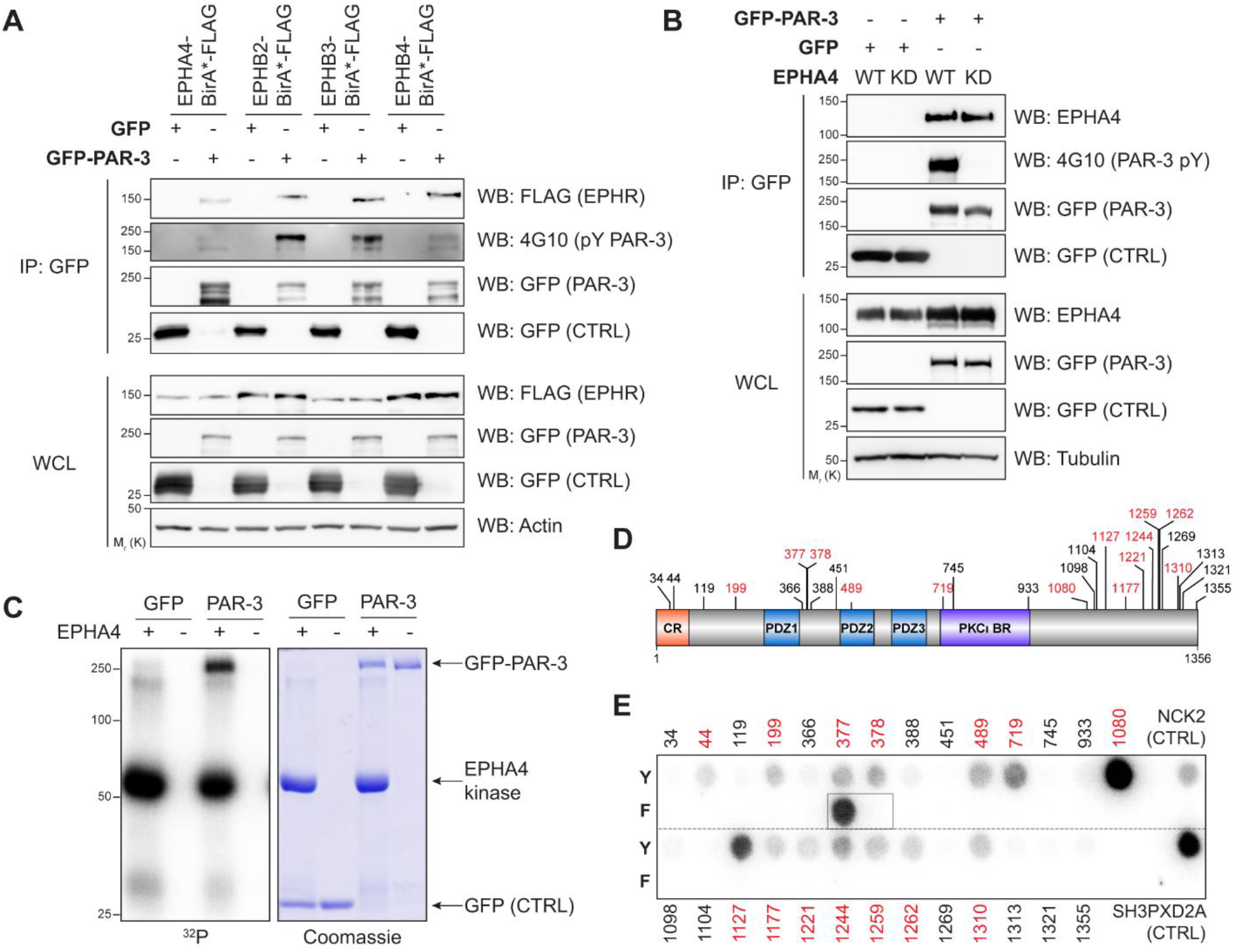
EPHA4 interacts with and phosphorylates PAR-3. (A) Western Blot analysis of EPHRs (FLAG) and PAR-3 Tyr phosphorylation (4G10) following co-transfection of GFP-PAR-3 and EPHR-BirA*-FLAG fusions and GFP affinity-purification in HEK293T cells. A representative image of 3 experiments is shown. (B) Western Blot analysis of EPHA4 and PAR-3 Tyr phosphorylation (4G10) following co-transfection of GFP-PAR-3 with wild-type (WT) or kinase-deficient (KD) EPHA4 in HEK293T cells. A representative image of 3 experiments is shown. (C) *In vitro* kinase assay using purified recombinant WT EPHA4 and GFP-PAR-3 affinity purified from HEK293T cells, in the presence of radioactive [γ-^32^P] ATP. Protein samples were resolved on SDS-PAGE, stained with Coomassie Brilliant Blue, and [γ-^32^P] incorporation was detected by autoradiography. A representative image of 2 experiments is shown. (D) Schematic of PAR-3 function domains and Tyr residues. CR: conserved region (oligomerization domain); PDZ: PSD95/Disc Large/ZO-1 domain; PKCι BR: PKCι (PRKCI) binding region. Peptides showing [γ-^32^P] incorporation >5 fold over their cognate non-phosphorylatable control (panel E) are highlighted in red. (E) Peptides corresponding to the 27 human PAR-3 Tyr and the corresponding non-phosphorylatable Phe counterpart were synthesized as 13-mer peptides and arrayed on a membrane. Phosphorylation was assessed by incubation with wild-type EPHA4 in the presence of radioactive [γ-^32^P] ATP, whose incorporation was detected by autoradiography. Peptide sequences and image quantification are provided in Table S4.

To examine PAR-3 phosphorylation by EPHA4, we performed an *in vitro* kinase assay using the recombinant EPHA4 catalytic domain (Dionne et al. 2018), and GFP-PAR-3 affinity purified from cell lysates as a substrate. We found that GFP-PAR-3 and not GFP control was phosphorylated by EPHA4 tyrosine kinase (Figure 5C). To pinpoint which of the 27 PAR-3 Tyr residues are targeted by EPHA4, we arrayed synthetic 13-mer peptides corresponding to each residue and surrounding sequences, as well as their non-phosphorylatable Phe counterpart. Using purified recombinant EPHA4, we performed a kinase assay with radiolabeled [γ-^32^P]. While 14 putative target Tyr were phosphorylated at levels above their cognate Phe control, we found that EPHA4 phosphorylated only two PAR-3 Tyr as strongly as the SH3PXD2A positive control (Dionne et al. 2018), namely Y1080 and Y1127 both positioned in PAR-3 C-terminus (Figures 5D and 5E, Table S4). This finding supports a direct phosphorylation of PAR-3 by EPHA4 on Tyr residues.

### EPHA4 phosphorylates PAR-3 Y1127 to create a docking site for C-terminal SRC kinase (CSK)

Protein phosphorylation can promote or preclude protein-protein interactions. To gain insight into the effect of PAR-3 phosphorylation on its interaction network, we performed affinity purifications of GFP-PAR-3 from HEK293T cells co-expressing WT or KD EPHA4, followed by MS identification. We completed experiments in triplicate, using GFP as a control bait to remove background via the SAINTexpress algorithm (Teo et al. 2014), with a BFDR threshold ≤1% (Figure S7, Table S2). Our analysis identified a total of 183 candidates associated with PAR-3 regardless of the presence of EPHA4 (Figures 6A and S7); 13 of them were already reported among the 78 PAR-3 interactors in the BioGRID database (17%) (Stark et al. 2006), including PAR6B, AMOT, PKCι (PRKCI) and YWHA isoforms. This also revealed 48 candidates that were absent and 37 candidates that were gained in the presence of EPHA4 WT or KD (Figures 6A-B and S6). This suggested that the presence of EPHA4 receptor modulates the PAR-3 interactome. Among PAR-3 interactors gained in cells expressing a catalytically active EPHA4, we found two proteins, GRB2 adaptor and C-terminal SRC kinase (CSK) that possess SRC-homology (SH) 2 domains recognizing pTyr residues (Figure 6B). PAR-3 does not have a binding consensus sequence for GRB2 SH2 domain (pY-X-N-X), thus indicating a putative indirect interaction. However, CSK SH2 was previously reported to directly bind PAR-3 Y1127 (Yang et al. 2009).

**Figure 6.**
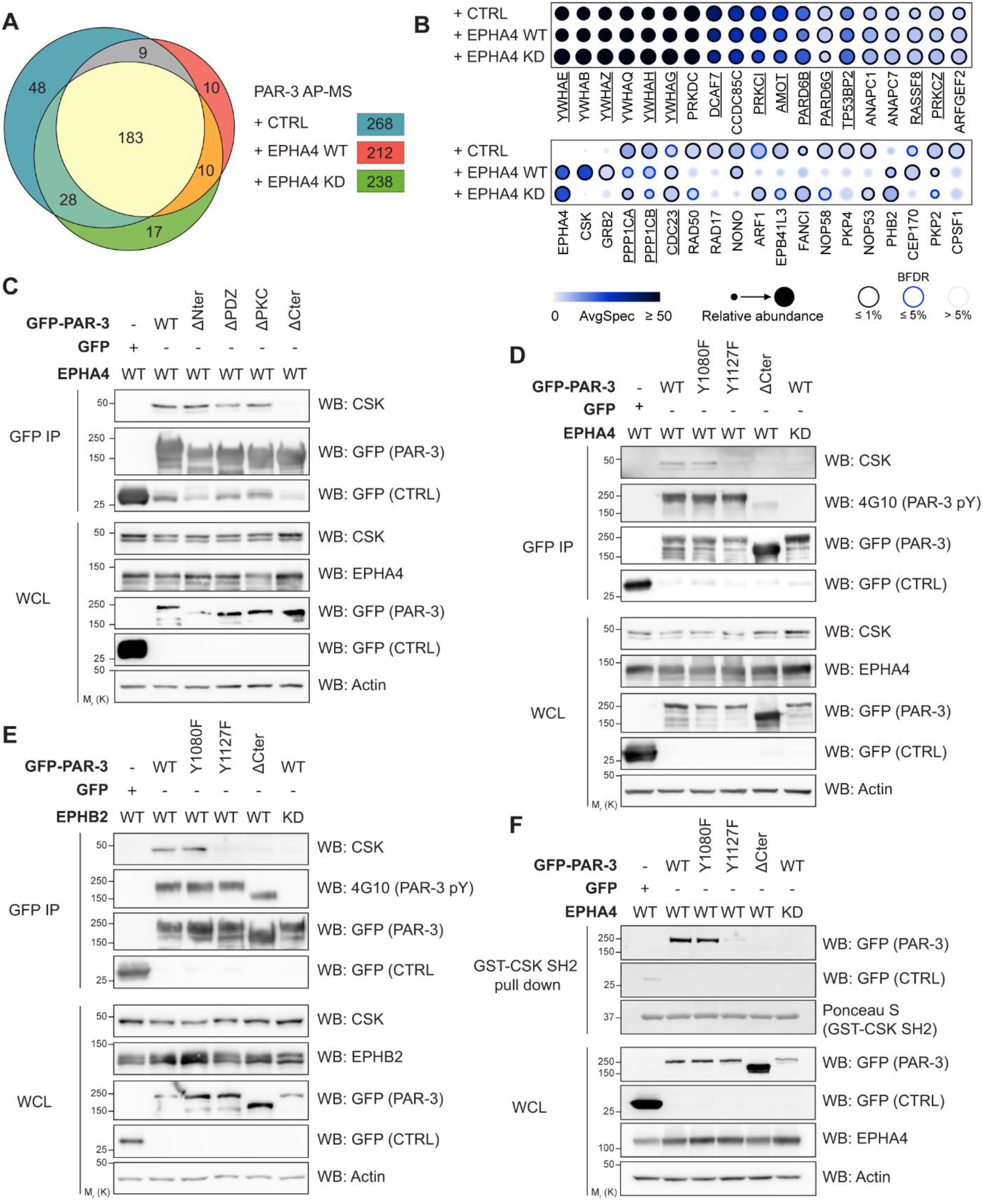
EPHA4 phosphorylates PAR-3 Y1127 to create a docking site for C-terminal SRC kinase (CSK). (A) Venn diagram representing the number of *bona fide* components of the PAR-3 interactome in the absence/presence of EPHA4 WT or KD, following the removal of background contaminants and non-specific interactions via SAINTexpress using a GFP control (BFDR <1%). (B) Dotplot representation of changes in abundance of a subset of identified PAR-3 interactors. Candidates previously reported in the BioGRID database are underlined. (C) Western Blot analysis of endogenous CSK following GFP affinity purification in HEK293T cells transfected with WT GFP-PAR-3 or domain deletion mutants, as well as WT EPHA4. A representative image of 3 experiments is shown. (D-E) Western Blot analysis of endogenous CSK and PAR-3 Tyr phosphorylation (4G10) following GFP affinity purification in HEK293T cells transfected with WT GFP-PAR-3 or mutants, as well as WT/KD EPHA4 (panel D) or EPHB2 (panel E). A representative image of 3 experiments is shown. (F) Western Blot analysis of GFP-PAR-3 and GFP following recombinant GST-CSK SH2 domain pull-down on lysates from cells transfected with WT GFP-PAR-3 or mutants, as well as WT or KD EPHA4 in HEK293T cells. A representative image of 3 experiments is shown.

We validated this association via GFP-PAR-3 affinity purification and endogenous CSK detection (Figure 6C), and tested which PAR-3 functional domains were required for the interaction (Figure 5D). We found that only PAR-3 C-terminus, which contains the most Tyr residues, was required (Figure 6C). To determine whether EPHA4-induced phosphorylation of PAR-3 Y1127 is sufficient to recruit CSK, we co-transfected WT or KD EPHA4 with WT, Y1080F, Y1127F or ΔCter GFP-tagged PAR-3. Following GFP-PAR-3 affinity purification, we detected endogenous CSK only in the presence of both WT EPHA4 and Y1127 (Figure 6D). We validated these findings with EPHB2 as well (Figure 6E), suggesting a conserved mechanism in the EPH family. Consistent with these observations, a recombinant GST-CSK SH2 domain used as a bait in a pull-down experiment with lysates expressing combinations of EPHA4 and PAR-3 only bound to the latter in the presence of a catalytically active EPHA4 and an intact Y1127 (Figure 6F). Together, these observations reveal that PAR-3 and CSK form a complex that is dependent on EPH-directed PAR-3 Y1127 phosphorylation.

To evaluate the requirement for CSK downstream of EPHR to establish cell segregation, we tested the effect of CSK loss-of function. We depleted CSK in EPH-expressing cells using two distinct siRNAs. Using our co-culture assay, we found that the average cluster size of CSK-depleted EPHB2 cells was significantly reduced compared to the control condition (p<0.0001 for both siRNAs; unpaired t-test) (Figure 7A-C). Collectively, our data reveal that PAR-3 and CSK form a complex that is dependent on EPH-directed PAR-3 Y1127 phosphorylation to control cell segregation (Figure 7D).

**Figure 7.**
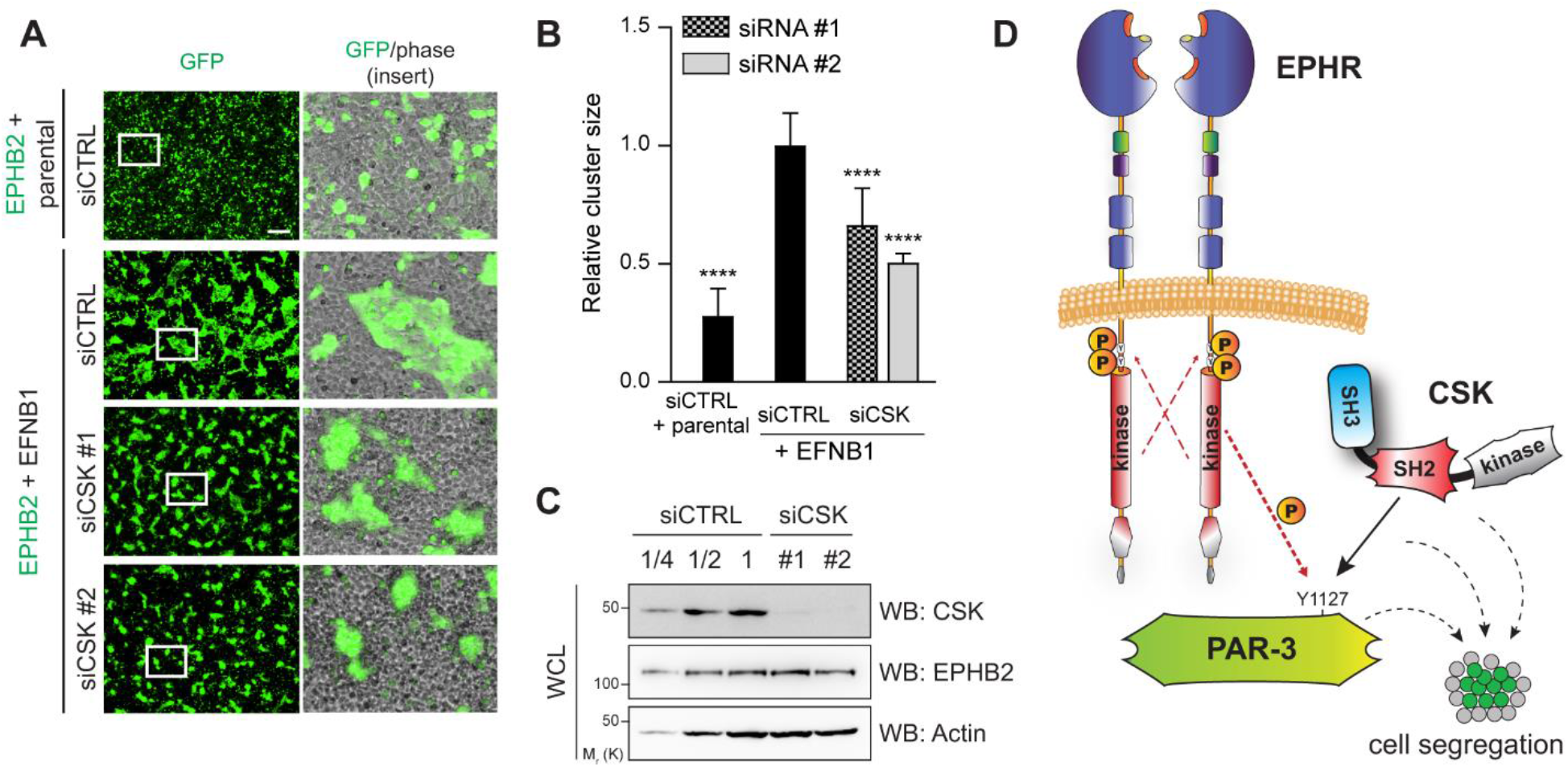
CSK depletion blocks EPHR-dependent cell segregation. (A) Representative images of cell co-cultures following siRNA depletion of CSK in EPHB2 expressing cells and mixing with ephrinB1-expressing cells, visualized by GFP (scale bar: 250 µm). (B) Quantification of the average size of EPHB2-GFP cell clusters depleted with two independent siRNAs targeting CSK, following mixing with ephrinB1-expressing cells, relative to control (EPHB2 siCTRL cells mixed with ephrinB1 cells). The mean of two experimental replicates is shown. Error bars indicate SD (**** p < 0.0001; unpaired t-test). (C) Western blot analysis of endogenous CSK following siRNA depletion. Successive dilutions are indicated to estimate depletion efficiency. A representative image from 2 experiments is shown. (D) Schematic representation of the proposed model for EPHR-PAR-3-CSK signalling complexes. EPHR associates with PAR-3 and directly phosphorylates PAR-3 on selected Tyr residues, including Y1127. This creates a docking site for CSK SH2 domain. The function of both PAR-3 and CSK downstream of EPHRs is required for cell segregation.

## DISCUSSION

We report a BioID-based high-confidence proximity network for EPHA4, EPHB2, EPHB3 and EPHB4. The identification of 386 associations that are not in the BioGRID database highly increase the set of reported interactions for EPHRs. It also stresses the complementarity between proximity labeling proteomics and other approaches based on the isolation of stable complexes, such as AP-MS. This is particularly striking for membrane proteins such as RTKs, due to the difficulty in solubilizing and maintaining intact the complexes during the purification process. Nonetheless, methods such as BioID have their own caveats, which were previously detailed (Gingras, Abe, and Raught 2019). For example, the residence time of potential preys in proximity to the bait will bias target identification. The selection of adequate controls, in particular when investigating membrane proteins, also represents a challenge.

We opted to use a YFP-BirA*-FLAG chimera that localized throughout our HEK293 cells (Figure 1B) as a control to remove non-specific interactions. We did consider using a membrane-targeted BirA* protein, for example via prenylation, as it initially appeared to be better-suited for our BioID experiments with EPHRs. However, proximity biotinylation MS experiments performed using a FLAG-BirA*-YFP-CAAX bait revealed that this control was too stringent, as it led to the identification of a large number of *bona fide* EPHR-associated candidates including AFDN, SHB, PIK3R1, YES1 and TIAM1 (data not shown). This was possibly due to the abundance and/or residence time of these preys at the plasma membrane. As a result, we reasoned that any target identified only with a subset of the 4 EPHRs tested would be deemed a *bona fide* proximity association, the remaining EPHR(s) acting as internal controls for specificity. For the 84 targets passing our SAINT threshold with all 4 EPHRs tested (i.e. “core” candidates), we surveyed background contaminants commonly identified in BioID experiments (CRAPome) (Mellacheruvu et al. 2013). We found that only 9/84 of the candidates were found in >10% of the experiments reported in the database, suggesting that they are not common contaminants but rather specific to EPHRs.

Our MS data highlighted nearly half (n=167) of the 395 proteins that were strictly associated with one of the four EPHRs used as bait (42%), while an equivalent number was common to at least three (39%) (Figures 2A and S2). These observations suggest the existence of both core signaling complexes that may be acting regardless of the EPHR that initiates signal, and of complexes specific to a given EPHR. While additional experiments are required to validate these observations, our data provide a strong basis to explore the molecular mechanisms underlying EPHRs signalling specificity via the selective recruitment of intracellular effectors. How in some instances EPHRs display overlapping functions and in others regulate specific processes is still poorly understood. This may be in part attributed to the recruitment of different combinations of downstream signaling effectors, as recently described for highly specific SAM–SAM heterotypic interactions (Wang et al. 2018). However, specific EPHR functions can be also achieved via distinct binding modes of a given effector protein to the same receptor, engaging separate EPHR intracellular domains (Zhuang et al. 2007, Ashlin et al. 2019). Therefore, further characterization of regulatory proteins that are common and/or specific to a given receptor would address the long-standing question of how EPHRs can transmit diverse, even opposite signals via their highly similar cytoplasmic domains.

Based on functional annotation with GO analysis, we determined that the identified proteins are involved in several cellular processes well established to be regulated by EPH signaling. This further increased our confidence that the BioID-identified candidates are putative EPHR effector proteins. Additionally, our analysis also associated EPHR BioID data with organization of cell-cell junctions. EPHA2 was recently reported to be localized at tight junctions and to be required for their organization via its association with AFDN in epidermal cells (Perez White et al. 2017). It is noteworthy that our BioID experiments detected AFDN in EPHA4, -B3 and -B4 proximity networks (Figure S2). The localization of EPHA2 at tight junctions in polarized epithelial cells is consistent with PAR-3 compartmentalization, suggesting that the kinase-substrate relationship may occur in different cell types, to regulate EPHR-dependent biological processes. For the two EPHRs that we found to be expressed in Caco-2 cells, EPHA1 and EPHB4, we describe compartmentalization in the basal and lateral domains, but not within tight junctions (Figure 3A). Altogether, the complementary distribution of EPHRs points to an involvement for EPHRs in the regulation of the different polarity complexes that are asymmetrically distributed in polarized epithelial cells (Laprise and Tepass 2011, Campanale, Sun, and Montell 2017). In addition, it is possible that alternative subsets of the 14 human EPHRs are expressed in different epithelial cell lines and tissues, as is the case in neural cells (Cramer and Miko 2016).

BioID proximity labeling proteomics experiments using EPHRs as baits were previously published. Perez White *et al*. first reported EPHA2 proximity networks of 131 and 215 proteins in 2D NHEK and 3D RHE cells, respectively (Perez White et al. 2017). Although they identified a few polarity proteins such as SCRIB and MARK2, cell polarity was not highlighted in their gene ontology classification. More recently, Lahaie *et al*. delineated the EPHB2 proximity network in HEK293 cells (Lahaie et al. 2019) and did find cell polarity, but not cell morphogenesis, among the best represented biological processes in our dataset. Finally, a BioID-based reference map of human HEK293 cells is being generated by the Gingras lab (cell-map.org) (Go et al. 2019). This endeavor includes proximity networks for six EPHRs as baits (i.e. EPHA2, -A3, -A4, -A7 and EPHB2, -B4). The most extensive dataset, which was obtained with EPHA2 as a bait, revealed several polarity proteins including DLG1, LLGL1, MARK2, EPB41L5, VANGL1 and SCRIB, which were also identified as common to all four EPHRs used as baits in our BioID experiments (Figures 2C and S2). Conversely, PARD3 (PAR-3) BioID unraveled EPHA2 and EFNB2 as preys. Not only does available BioID data suggest a link between EPHRs and cell polarity, but also a function for PAR-3 in EPHR signaling, which we further supported in this report.

We describe that PAR-3 is a substrate for EPHA4. A number of kinases were previously reported to modulate PAR-3 phosphorylation and function. Notably, serine/threonine kinases PKCι (Nagai-Tamai et al. 2002), Par1 (MARK2) (Benton and St Johnston 2003), Aurora A/B (Khazaei and Püschel 2009), ERK2 (Funahashi et al. 2013, Yang et al. 2012), ROCK (Nakayama et al. 2008) and NDR1/2 (Yang et al. 2014) were all described to regulate PAR-3 protein interactions and function, in some cases via a dynamic interplay involving protein phosphatase PP1 (Traweger et al. 2008). Although multiple PAR-3 Tyr residues were also discovered to be phosphorylated following cell stimulation by RTK ligands EGF (Wang et al. 2006), HGF (Organ et al. 2011) and EFNB1 (Jørgensen et al. 2009), none of their cognate RTKs were described to directly phosphorylate PAR-3. For EGF/EGFR-induced PAR-3 phosphorylation, the authors found that Y1127, Y1177 and Y1321 were targeted via a mechanism that involves the direct phosphorylation of PAR-3 by c-SRC (Wang et al. 2006). In addition, they demonstrated that Y1127 phosphorylation impaired the association between PAR-3 and LIM kinase 2 (LIMK2), a step that is required for adequate epithelial tight junction assembly. Phosphorylation of PAR-3 Y1127 was also reported to create a novel pTyr binding motif for the c-SRC negative regulator, CSK, via its unique SH2 domain (Yang et al. 2009). We found that EPHA4 associates with and directly phosphorylates PAR-3 on Y1080 and Y1127, and that the latter is a docking site for CSK. This significantly adds to previous findings (Wang et al. 2006, Yang et al. 2009). Remarkably, we uncovered that the PAR-3-CSK cassette is required to regulate EPHR-dependent cell segregation. To our knowledge, these findings provide the only example to date of an RTK acting as a PAR-3 kinase. Collectively, our data indicate that PAR-3 Y1127 is a major target for PAR-3 regulation and is crucial for its function as a scaffold for the assembly of signalling networks.

Although our BioID MS identified a few hundreds putatively novel EPHR associated proteins, the experimental setup required for BirA* to achieve sufficient biotinylation of endogenous proteins did not allow to obtain time-resolved stimulation following ligand application, but rather provided a global picture of EPHRs proximity networks regardless of the activation state. Interestingly, we identified receptor activation-dependent candidates such as NCK2 with EPHA4 (Figure S2), an association that is dependent on phosphorylation of Y596/602 on the receptor (Dionne et al. 2018). This indicates that either EPHRs display a baseline activation level despite not being stimulated with an exogenous cognate ligand in cultured HEK293 cells, or that EPHR activation-dependent candidates may also associate in an alternative manner. Notwithstanding, the development of more potent BirA*-derived biotin ligases such as TurboID, which achieves significant biotinylation of endogenous proteins within 10-30 minutes (Branon et al. 2018), should allow probing for time-resolved, ligand-specific EPHR proximity networks.

## ACKNOWLEDGMENTS

We thank members of the Bisson laboratory for discussions, A. Freywald and L. McCaffrey for critical reading of the manuscript, F.J.M. Chartier and U. Dionne for expert technical assistance, as well as A. Cailler, C. Dziengielewski, L. McCaffrey and L.T. Wang for constructs and reagents. Mass spectrometry was performed at the CHU de Québec – Université Laval Proteomics Platform. This work was funded by a FRQNT Team Grant (2019-PR-254641) to N.B., S.E. and P.L.. N.B. was also supported by funds from the Canada Foundation for Innovation (30308, 34963) and holds a Canada Research Chair (Tier 2) in Cancer Proteomics. S.B. held a PROTEO scholarship during the completion of this work. N.L. holds an Alexander-Graham-Bell Canada Graduate Scholarship from NSERC, and previously a FRQS Doctoral Award and a Frederick-Banting and Charles-Best Canada Graduate Scholarship from CIHR.

## AUTHOR CONTRIBUTIONS

Investigation: S.L.B., N.L., K.J., F.L., J.P.L.; Resources: J.N.L., S.E., P.L., N.B.; Writing – Original Draft: S.L.B., N.B.; Writing – Review & Editing: all authors; Visualization: S.L.B., N.L., K.J., F.L.; Supervision: J.N.L., S.E., P.L., N.B.; Project Administration: N.B.; Funding Acquisition: S.E., P.L., N.B.

## STAR METHODS

### CONTACT FOR REAGENT AND RESOURCE SHARING

Further information and requests for resources and reagents should be directed to and will be fulfilled by the Lead Contact, Nicolas Bisson.

### EXPERIMENTAL MODEL DETAILS

#### Cell culture, transfections and transductions

Human embryonic kidney 293 (HEK293) Flp-In T-Rex, 293T cells (HEK293T) and Caco-2 human colorectal adenocarcinoma cells were cultured in Dulbecco’s Modified Eagle’s medium (DMEM, Thermo Fisher Scientific) high glucose supplemented with 10% fetal bovine serum (Sigma-Aldrich) at 37°C under 5% CO2. Stable Flp-In T-REx HEK293 lines were generated as described (Kean, Couzens, and Gingras 2012), and pools of stable transfectants were selected with 100 μg.ml^-1^ hygromycin (Thermo Fisher Scientific). Expression was induced with 1 μg.ml^-1^ tetracycline for 24 h. For BioID experiments, cells were treated with 50 µM biotin for an additional 24h. For cell segregation experiments, stable HEK293 T-REx cells expressing WT EPHB2 and GFP, herein referred as HEK-EPHB2 & GFP cell line, were generated by co-transfection of pcDNA5-FRT-TO-EPHB2, pOG44 and pEGFP-C1 vectors followed by a 14-day co-selection with 100 μg.ml^-1^ hygromycin. Individual GFP positive clones were picked, expanded and analyzed for EPHB2 expression. HEK293T cells stably expressing shPARD3, as described in (Dziengelewski et al. 2020), were grown in the presence of 1.5 mg.ml^-1^ puromycin. Transient transfections were performed using polyethylenimine (PEI). Reverse transfections of siRNAs were performed using RNAiMAX (Thermo Fisher Scientific). Lentiviruses were produced by co-transfection of pLKO constructs, pMD2G (Addgene plasmid #12259, from Trono lab) and psPAX2 (Addgene plasmid #12260, from Trono lab). Viral supernatants were collected after 36h, filtered through a 0.45 µm membrane and supplemented with 8 μg.ml^-1^ polybrene (Sigma-Aldrich). Cells were transduced for 24h or 48h prior to experiments. Ephrin stimulation of EPH-expressing HEK293 cells was performed as described (Jørgensen et al. 2009).

### METHOD DETAILS

#### Constructs and sequences

For stable expression in mammalian cells, human sequences of EPHA4, EPHB2, EPHB3, EPHB4 were cloned into pcDNA5-FRT-TO-BirA*-FLAG or pcDNA5-FRT-TO vectors (Thermo Fisher Scientific) and human EFNB1 and EFNB2 were subcloned into pcDNA5-FRT-TO with a C-terminal 2xHA tag. Previously described murine EPHA4 WT and KD in pcDNA3.1(-) (Dionne et al. 2018), rat PARD3 cDNA in pEGFP-C1 (Hidalgo-Carcedo et al. 2011), and PARD3 domain deletion mutants (Dziengelewski et al. 2020) were used for transient expression. Human sequences of EPHA1, EPHA2, EphA4 and EphB4 were cloned into pLJM1-EGFP vector (Addgene plasmid # 19319, from David Sabatini) for transient infections. Point mutations were introduced using the Q5 mutagenesis kit (New England Biolabs) and all inserts were verified by sequencing. Dicer-Substrate Short Interfering RNAs (DsiRNAs) sequences (Integrated DNA Technologies) are provided (supplemental Table S3). Control non-targeting DsiRNA sequences were provided by the manufacturer (Integrated DNA Technologies). ShRNA sequence targeting human PARD3 (5’-CCG GGC CAT CGA CAA ATC TTA TGA TCT CGA GAT CAT AAG ATT TGT CGA TGG CTT TTT G-3’), PKCι (TRCN0000219727), EPHA1 (TRCN0000314913 and TRCN0000314914) and EPHB4 (TRCN0000314827 and TRCN0000001773) were inserted downstream of the U6 promoter in the pLKO vector (Addgene plasmid #8453, from Trono lab). Control vector was pLKO with a non-targeting shRNA sequence designed in Stephane Gobeil’s lab (5’-CCG GTC CTA AGG TTA AGT CGC CCT CGC TCG AGC GAG GGC GAC TTA ACC TTA GGT TTT T-3’). EGFP sequence was inserted in all pLKO vectors to monitor the rate of infected cells. Vectors used for the bacterial expression of GST-tagged recombinant juxtamembrane and catalytic domains of EPHA4, and SH2 domain of CSK were previously described (Dionne et al. 2018, Huang et al. 2008).

#### Cell lysis and affinity purification

BioID experiments were performed as previously described (Jacquet et al. 2018). Briefly, cells were lysed in RIPA buffer (50 mM tris-HCl pH 7.5, 150 mM NaCl, 1 mM EDTA, 1 mM EGTA, 1% NP-40, 0.1% SDS, 0.5% sodium deoxycholate) supplemented with protease inhibitors (1 mM PMSF, 10 μg.ml^-1^ aprotinin, 10 μg.ml^-1^ leupeptin,10 μg.ml^-1^ pepstatin) and 250 U of benzonase. Following 1h incubation with agitation, lysates were sonicated, centrifuged 30 min at 20 000 g and cleared supernatants were further incubated with streptavidin agarose beads (Sigma-Aldrich) for 3h with agitation at 4°C. For FLAG and GFP affinity purifications, as well as Western blotting, cells were lysed in KLB buffer (20 mM Tris-HCl pH 7.4, 150 mM NaCl, 1 mM EDTA, 0.5% sodium deoxycholate) supplemented with protease inhibitors. Phosphatase inhibitors (Cocktail 2, Sigma-Aldrich) were added if required. Lysates were incubated 15 minutes on ice with occasional gentle agitation followed by 20 min centrifugation at 20 000 g. Protein concentrations were measured and normalized using BCA kit (Thermo Fisher Scientific). Cleared supernatants were incubated with FLAG-M2 agarose beads (Sigma-Aldrich) or GFP-Trap agarose beads (Chromotek) for 2h at 4°C with rotation. For mass spectrometry, beads were washed three times with lysis buffer, followed by two washes with 20 mM Tris-HCl (pH 7.4). Proteins were eluted with 50 mM phosphoric acid, three times for 10min and stored at −80°C. Eluted proteins were digested with Promega Sequencing Grade Modified Trypsin (Thermo Fisher Scientific) as described (Beigbeder et al. 2016). The resulting peptides were desalted using StageTips (Rappsilber, Mann, and Ishihama 2007) and dried down by vacuum centrifugation prior to LC-MS/MS analyses. For Western blotting, beads were washed three times with lysis buffer then boiled in 4x Laemmli buffer (Laemmli 1970).

#### Mass spectrometry

Dried peptides were resuspended in 15 µl of loading solvent (2% acetonitrile, 0.05% TFA), and 5 µl was used for injection onto a 300 µm inner diameter x 5 mm C-18 Pepmap cartridge precolumn (Dionex / Thermo Fisher Scientific) at 20 µl/min in loading solvent. After 5 min of desalting, the precolumn was switched online with a 75 μm inner diameter x 50 cm separation column packed with 3 μm ReproSil-Pur C18-AQ resin (Dr. Maisch HPLC GmbH) equilibrated in 95% solvent A (2% acetonitrile, 0.1% formic acid) and 5% solvent B (80% acetonitrile, 0.1% formic acid). Peptides were separated and eluted over a 90 min gradient of 5% to 40% solvent B at 300 nl/min flow rate generated by an UltiMate(tm) 3000 RSLCnano system (Dionex / Thermo Fisher Scientific) and analyzed on an Orbitrap Fusion mass spectrometer equipped with a nanoelectrospray ion source (Thermo Fisher Scientific). The Orbitrap Fusion was operated in data-dependent acquisition mode with the XCalibur software version 3.0.63 (Thermo Fisher Scientific). Survey MS scans were acquired in the Orbitrap on the 350 to 1800 m/z range using an automatic gain control (AGC) target of 4e5, a maximum injection time of 50 ms and a resolution of 120,000. The most intense ions per each survey scan were isolated using the quadrupole analyzer in a window of 1.6 *m*/*z* and selected for Higher energy Collision-induced Dissociation (HCD) fragmentation with 35% collision energy. The resulting fragments were detected by the linear ion trap at a rapid scan rate with an AGC target of 1e4 and a maximum injection time of 50ms. Dynamic exclusion was employed within a period of 20 s and a tolerance of 10 ppm to prevent selection of previously fragmented peptides. All MS/MS peak lists were generated using Thermo Proteome Discoverer version 2.1 (Thermo Fisher Scientific).

#### Data-Dependent Acquisition MS analysis

Mass spectrometry data was stored, searched and analyzed using the ProHits laboratory information management system (LIMS) platform (Liu et al. 2016). RAW MS files were converted to mzML and mzXML using ProteoWizard (3.0.4468; (Kessner et al. 2008)). The mzML and mzXML files were then searched using Mascot (v2.3.02) and Comet (v2012.02 rev.0). Spectra were searched with the RefSeq database (version 57, January 30th, 2013) acquired from NCBI against a total of 72,482 human and adenovirus sequences supplemented with “common contaminants” from the Max Planck Institute (http://141.61.102.106:8080/share.cgi?ssid=0f2gfuB) and the Global Proteome Machine (GPM; http://www.thegpm.org/crap/index.html). Charges +2, +3 and +4 were allowed and the parent mass tolerance was set at 12 ppm while the fragment bin tolerance was set at 0.6 amu. Carbamidomethylation of cysteine was set as a fixed modification. Deamidated asparagine and glutamine and oxidized methionine were allowed as variable modifications. The results from each search engine were analyzed through TPP (the Trans-Proteomic Pipeline (v4.6 OCCUPY rev 3) (Deutsch et al. 2015), via the iProphet pipeline (Shteynberg et al. 2011).

#### Statistical rationale and MS data analysis with SAINTexpress

Each BioID and AP-MS experiment was performed in biological triplicate and was processed independently on different days using cells from successive passages. Controls for each experiment were treated concomitantly to experimental samples. YFP-BirA*-FLAG was used as a control for EPHR-BirA*-FLAG in BioID experiments and GFP was used as a control for GFP-PARD3 in AP-MS experiments. To minimize carry-over issues during liquid chromatography, extensive washes were performed between each sample. Sample loading order was randomized. MS data was analyzed with SAINTexpress, a simplified version of the Significance Analysis of INTeractome method (Teo et al. 2014), to distinguish background contaminants and non-specific interactions from *bona fide* protein associations. The probability value of each potential protein-protein interaction compared to background contaminants was calculated via SAINTexpress version 3.6.1 (Teo et al. 2014), using default parameters. The three control samples were used in uncompressed mode. Two unique peptide ions and a minimum iProphet probability of 0.95 were required for protein identification prior to SAINTexpress. Complete lists of SAINTexpress analyses results are provided in (supplemental Tables S1 and S2). Decoy and common contaminant proteins were removed as indicated in Table S1. Interaction networks with SAINT outputs with BFDR ≤ 0.01 were generated using Cytoscape (version: 3.7.2) (Shannon et al. 2003). ClueGO plug-in v2.1.7 set for GO Biological process analysis with upper medium network specificity was used for clustering analysis (Bindea et al. 2009). Changes in interactomes were visualized using DotPlot, a protein-protein interaction data visualization tool (Knight et al. 2017). Venn diagrams were made using Venn Diagram Plotter (Pacific Northwest National Laboratory, https://omics.pnl.gov/software).

#### Data availability

The mass spectrometry proteomics data have been deposited to the ProteomeXchange Consortium (ProteomeXchange ID: PXD021257) via the MassIVE partner repository (http://massive.ucsd.edu), with the dataset identifier MSV000086054. with the password “polarity” to access files prior to publication. Additional files include a list of BioID and AP-MS experiments, controls, and sample files, the complete SAINTexpress outputs for each dataset as well as a ‘‘README’’ file that describes the experimental procedures.

#### Western blotting

Proteins were separated by SDS-PAGE and transferred onto nitrocellulose membranes. Immunoblot analysis was performed as described before (Jacquet et al. 2018). Primary antibodies used were: rabbit anti-FLAG (Sigma-Aldrich, #F7425), mouse anti-pan phosphotyrosine 4G10 (Millipore, #05-321), mouse anti-GFP (Abcam, #290), mouse anti-EPHA4 (BD Biosciences, #610471), goat anti-EPHB2 (R&D Systems, # AF467), goat anti-EPHA1 (R&D Systems, #AF638), mouse anti-EPHA2 (Millipore, #05-480), goat anti-EPHB4 (R&D Systems, #AF3038), mouse anti-PKCι (BD, #610175), rabbit anti-CSK (Cell Signaling Technology, #4980), rabbit anti-PAR-3 (Millipore, #07-330), mouse anti-actin (Cell Signaling Technology, #3700). Secondary antibodies were the following: horse anti-mouse IgG HRP (Cell Signaling Technology, #7076), goat anti-rabbit IgG HRP (Cell Signaling Technology, #7074), rabbit anti-goat IgG HRP (Thermo Fisher Scientific, #31402), rat anti-mouse TrueBlot HRP (Rockland, #18-8817-30), goat anti-mouse IgG Alexa Fluor 647 (Thermo Fisher Scientific, #A-21235), goat anti-rabbit IgG Alexa Fluor 647 (Thermo Fisher Scientific, #A-21245). Biotinylated proteins were detected using streptavidin-HRP conjugate (Thermo Fisher Scientific, #434323). Signal was revealed using Clarity western ECL substrate (Bio-Rad) and acquired on Amersham Biosciences Imager 600RGB (GE Healthcare).

#### Immunofluorescence

HEK293 T-Rex cells were pre-fixed with 2% paraformaldehyde (BioShop) for 5 min, washed with PBS and fixed with 4% paraformaldehyde for 15 min at room temperature. Cells were permeabilized with 0.2% Triton X-100 (Sigma-Aldrich) for 15 min, blocked in 10% goat serum (Wisent) with 0.1% NP-40 (Sigma-Aldrich) in PBS and incubated with rabbit anti-FLAG (Sigma-Aldrich, #F7425) primary antibody diluted in blocking solution for 1h at room temperature. Following 3 washes in PBS with 0.1% NP-40, cells were incubated for 1h with Alexa Fluor 488 goat anti-rabbit IgG (Jackson ImmunoResearch, #111-545-003) secondary antibody. Cells were washed 3 times with 0.1% NP-40 in PBS and nuclei were stained with DAPI (Sigma-Aldrich) for 5 min. Slides were washed twice with PBS and mounted in ProLong Gold Antifade (Thermo Scientific). For Caco-2 spheroids, cells were fixed after six days of culture on Matrigel with 4% paraformaldehyde for 15-60 min at room temperature or 37°C. Cells were washed 3 times for 15 min with PBS, then permeabilized and blocked for 1 h at room temperature in 0.5% Triton X-100 (Sigma-Aldrich), 10% donkey serum (Sigma) in PBS. Cells were incubated with goat anti-EPHA1 (R&D, #AF638), goat anti-EPHB4 (R&D, #AF3038), rabbit anti-Ezrin (Cell signaling, #3145S) and mouse anti-ZO-1 (Thermo Fisher Scientific, #33-9100) primary antibodies overnight at 4°C. Following 3 washes of 15 min in blocking solution, cells were incubated for 2h with Alexa Fluor 488 donkey anti-goat IgG (Thermo Fisher Scientific, #A11055), Alexa Fluor 568 donkey anti-mouse IgG (Thermo Fisher Scientific, #A10037), Alexa Fluor 647 donkey anti-rabbit IgG (ThermoFisher Scientific, #A31573) and/or DAPI (Sigma-Aldrich). Cells were washed 3 times for 15 min in PBS and left in PBS until image acquisition. Images were acquired on a confocal laser scanning microscope (Olympus FV1000) using PlanApo N 60x Oil (numerical aperture 1.42) or PlanApo 40x (numerical aperture 0.90) objectives. Images were processed with Olympus Fluoview 3.1 B or ImageJ softwares.

#### Caco-2 spheroids formation

To produce Caco-2 cysts, single cells were plated on top of a 20 μL pre-coat of 80% Matrigel Growth Factor Reduced (Fisher Scientific, CB-40230A) and 20% Collagen I, Bovine (Fisher, CB-4023) in a µ-Slide 8 Well ibidi plate (ibidi, 80826). Trypsinized cells were resuspended in culture medium supplemented with 2% Matrigel Growth Factor Reduced; 10,000 cells were seeded per well. Cells were grown for six days before fixation for immunofluorescence. For loss-of-function experiments, cells were infected 24h prior to seeding on Matrigel. For Western blot analyses, 2D cultures were grown in parallel for six days.

#### Cell segregation

Cell segregation assay was previously described (Wu et al. 2019) and was performed with minor modifications. Briefly, HEK293 parental cells, HEK-EPHB2 & GFP cells and HEK-ephrinB1 cells were induced with tetracycline for 24h. Cells were dissociated in Enzyme-free cell dissociation buffer (Thermo Fisher Scientific), thoroughly resuspended in culture media supplemented with tetracycline and counted on an automated cell counter (Bio-Rad TC20). Homogenous cell populations were mixed 1:1, plated on µ-Dish 35 mm, high, ibiTreat (ibidi) pre-coated with 10 μg.ml^-1^ fibronectin (Sigma-Aldrich) at a final density of 0.4×10^6^ cells/cm^2^ and incubated for 48h (until confluence is reached). Cells were fixed with 4% paraformaldehyde for 15 min and left in PBS until imaging. For loss-of function studies, HEK-EPHB2 & GFP cells were either transfected with DsiRNA or transduced with viruses for 48h prior to cell mixing. Transmitted light and GFP images were acquired on Nikon Ti Eclipse motorized inverted microscope at room temperature using a Hamamatsu ORCA-ER 1344 × 1024 CCD camera. For each condition within a biological replicate, five independent and randomly selected fields were acquired using NIS Elements AR 5.02.00 acquisition software. Each field was composed of four images (2×2) acquired with a Plan Fluor 4x (numerical aperture 0.13) objective that were stitched with a 15% overlap via NIS Elements AR software, using GFP as a reference. Images were exported as a TIFF and five stitched GFP images were further used for cell segregation data analysis and quantification for each condition.

#### Image analysis of cell segregation

All TIFF images were systematically processed and analyzed using ImageJ (version 1.52n). First, a threshold was applied to each 16-bit image using a threshold function adjusted with the LI algorithm. Next, each particle size was retrieved by the ‘Analyze Particles’ function set to detect particles with a size (pixel^2) above 50 pixels and a circularity between 0.00-1.00. Outlines were also displayed and the particle map was compared and validated toward the original image. The average particle size was obtained based on all the particle sizes within the image. The average cluster size of HEK-EPHB2 & GFP cells from different conditions was normalized toward the control condition (HEK-EPHB2 & GFP siCTRL cells mixed with HEK-ephrinB1 cells) and represents two independent biological replicates. Statistical unpaired Student’s t-tests were performed with GraphPad Prism 8.

#### Protein expression and purification

Expression of GST-EPHA4 kinase (591-896) or GST-CSK SH2 domain was induced in BL21 (DE3) *E. coli* with 0.5 mM IPTG for 16h at 16°C. Bacteria were pelleted and resuspended in GST buffer (20 mM Tris-HCl pH 7.5, 0.5 M NaCl, 1 mM EDTA and 1 mM DTT) supplemented with protease inhibitors (P8340, Sigma) and lysed by sonication. Recombinant proteins were purified using GST beads (GE Healthcare), eluted with 30 mM GSH (Bio-Basic) and concentrated using Amicon 10K filters (Millipore). Protein concentration was determined toward known amounts of BSA on SDS-PAGE gels stained with Coomassie Brilliant Blue and acquired on Amersham Biosciences Imager 600RGB (GE Healthcare) and quantified with the Amersham Imager 600 analysis software (GE Healthcare).

#### Kinase assays

GFP-PAR-3 or GFP control were transiently overexpressed in shPAR-3 HEK293T cells and affinity purified on GFP-Trap agarose beads (Chromotek). Beads were washed with kinase buffer (10 mM MgCl2, 20 mM MnCl2, 2.5 mM DTT in 50 mM HEPES pH 7.5) supplemented with protease inhibitors (P8340, Sigma-Aldrich) and split into two equal parts. Beads were resuspended in kinase buffer containing 4 µCi of radiolabeled [γ-^32^P] ATP and 75 µM of non-labeled ATP, and then incubated in the presence or absence of 5 µg of recombinant EPHA4 for 1h at 37°C with 1100 rpm agitation. Beads were boiled in 4x Laemmli buffer to stop the reaction and protein samples were resolved by SDS-PAGE. Gels were stained with Coomassie Brilliant Blue to assess the loading consistency across the samples and then vacuum dried. For peptide arrays, peptides corresponding to the peptide environment of each of 27 Tyr residues on human PARD3, were synthesized as 13-mers with the Tyr residue in the middle position. For non-phosphorylatable controls, the Tyr was replaced by a Phe. Arrays were prepared as previously described (Dionne et al. 2018). Peptides sequences are listed in supplemental Table S4. The radioactive peptide array experiment was performed as described (Warner et al., 2008). For both assays, incorporation of radiolabeled [γ-^32^P] ATP was determined using a Fujifilm FLA-5100 imager. Signal quantification was performed using ImageJ software with Protein Array Analyzer macro (Gilles Carpentier; http://image.bio.methods.free.fr/ImageJ/?Protein-Array-Analyzer-for-ImageJ.html#outil_sommaire_0)

#### GST pull-down assay

WT or KD EPHA4 was co-transfected with WT, Y1080F, Y1127F or ΔCter GFP-tagged PAR-3 or GFP control into 293T cells stably expressing shRNA targeting endogenous PAR-3. Following the 36h of transient overexpression, cells were lysed in ice cold KLB buffer (20 mM Tris-HCl pH 7.4, 150 mM NaCl, 1 mM EDTA, 0.5% sodium deoxycholate) supplemented with protease (P8340, Sigma-Aldrich) and phosphatase inhibitors (Cocktail 2, Sigma-Aldrich). Lysates were incubated 15 minutes on ice, centrifuged for 20 min at 20,000g and protein concentration was determined using a BCA kit (Thermo Fisher Scientific). Supernatants were pre-cleared with glutathione beads for 2h at 4°C with rotation; beads were discarded. A total of 5 µg of recombinant GST-CSK SH2 domain or GST control coupled to glutathione beads was added to 1 mg of pre-cleared cell lysate for each pull-down analysis and incubated for 2h at 4°C with rotation. Beads were washed three times with wash solution (20 mM Tris-HCl pH 8.0, 200 mM NaCl, 1 mM EDTA, 0.5% NP-40) and then boiled in 4x Laemmli buffer and stored at −20°C for further Western blotting analysis.

## SUPPLEMENTAL FIGURES

**Figure S1.**
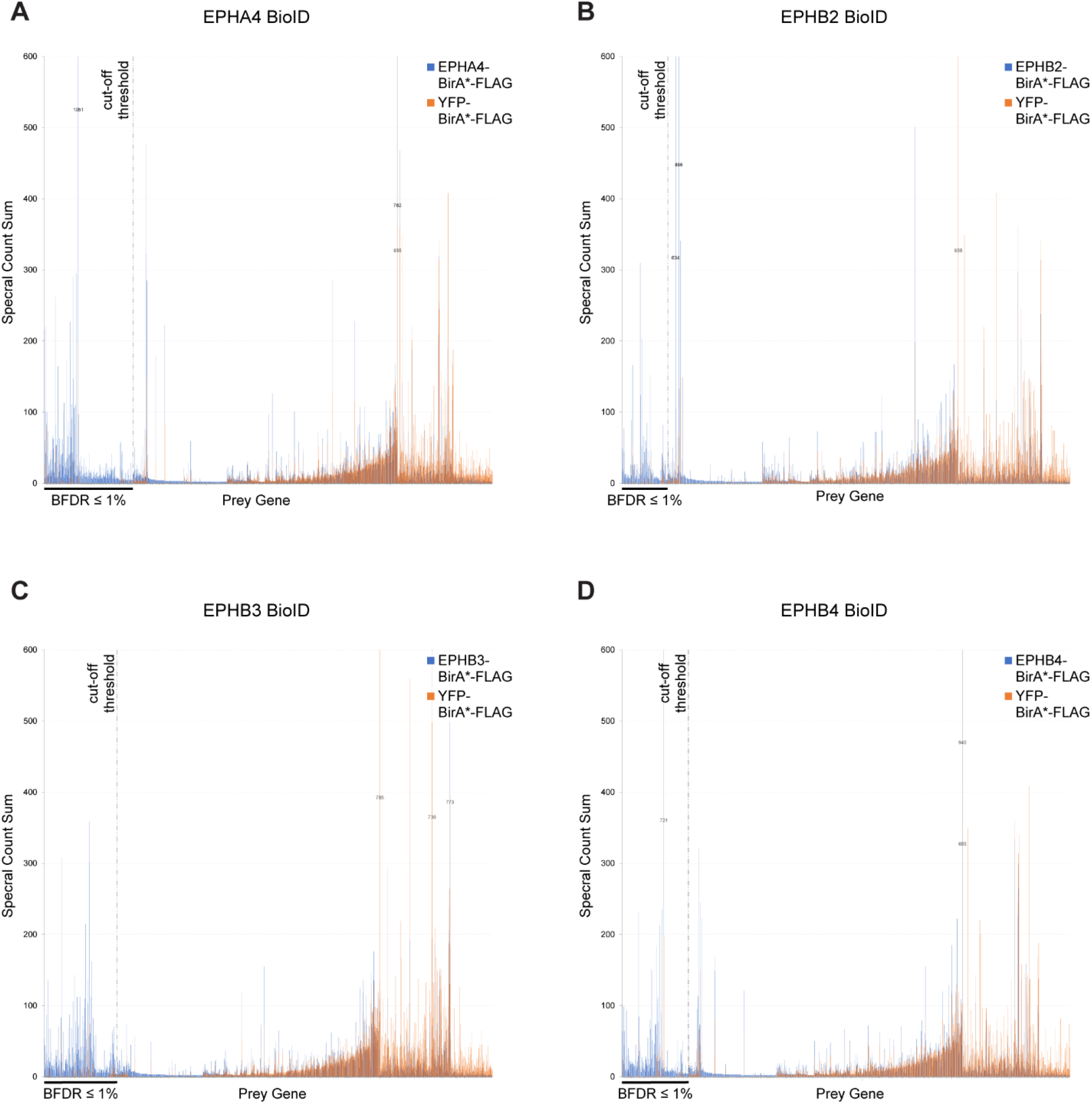
Candidate ranking for each EPHR BioID experiment. Spectral counts sum across the three experimental replicates for proteins identified in BioID experiments from cells stably expressing EPHR (blue) or control (orange) baits. A cutoff threshold corresponding to BFDR ≤1% following SAINTexpress analysis is highlighted with a dashed vertical line. A sum of spectral counts > 600 is indicated for specific preys whose intensity is beyond the graph scale.

**Figure S2.**
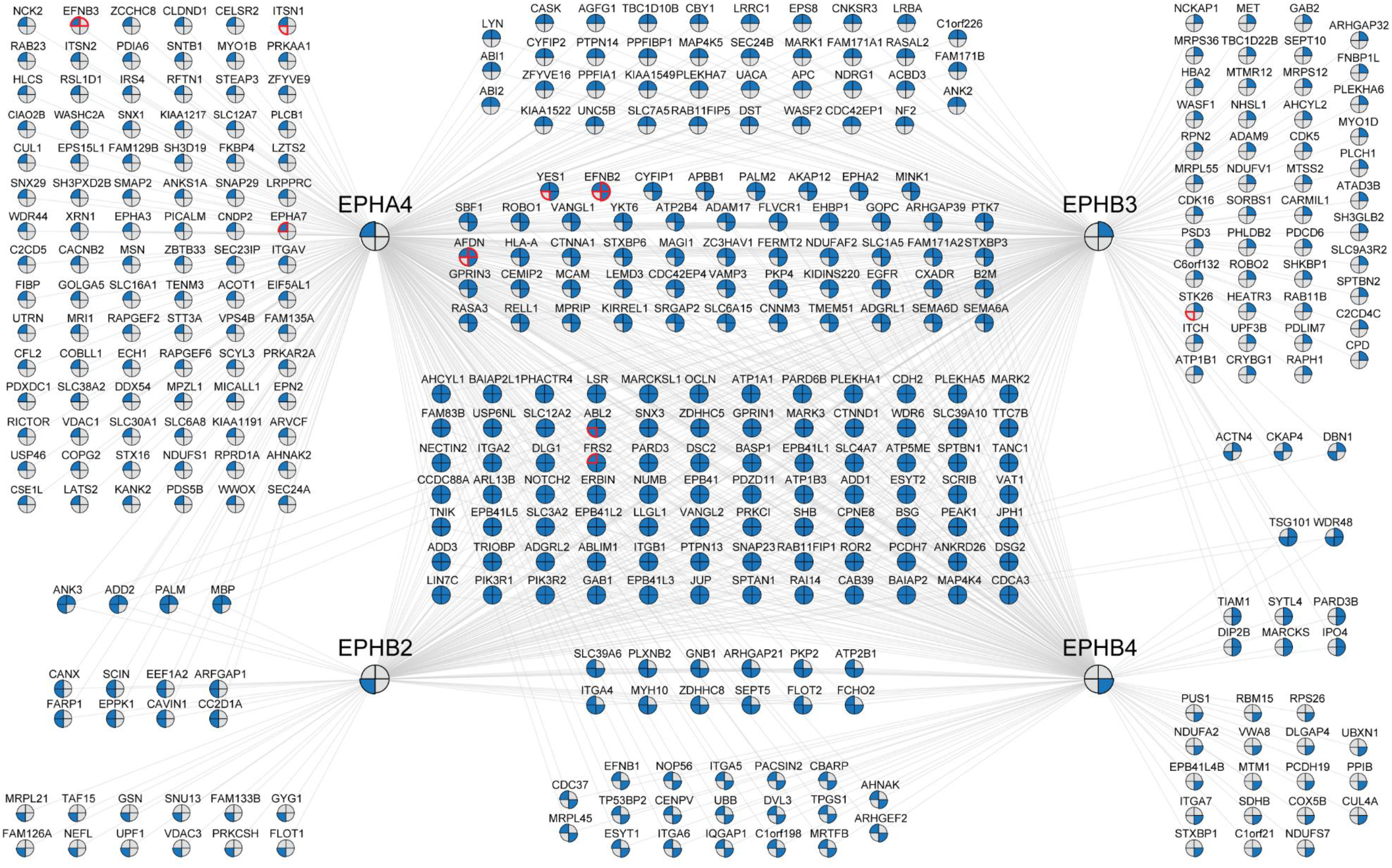
BioID proteomics identifies EPHA4, -B2, -B3, -B4 proximity networks. *Bona fide* components of EPHA4, EPHB2, EPHB3 and EPHB4 proximity networks following the removal of background contaminants and non-specific interactions via SAINTexpress using a YFP-BirA*-FLAG control (BFDR ≤1%). BioID experiments were performed in triplicate. Candidates previously reported in the BioGRID database for each EPHR bait are noted with a red edge.

**Figure S3.**
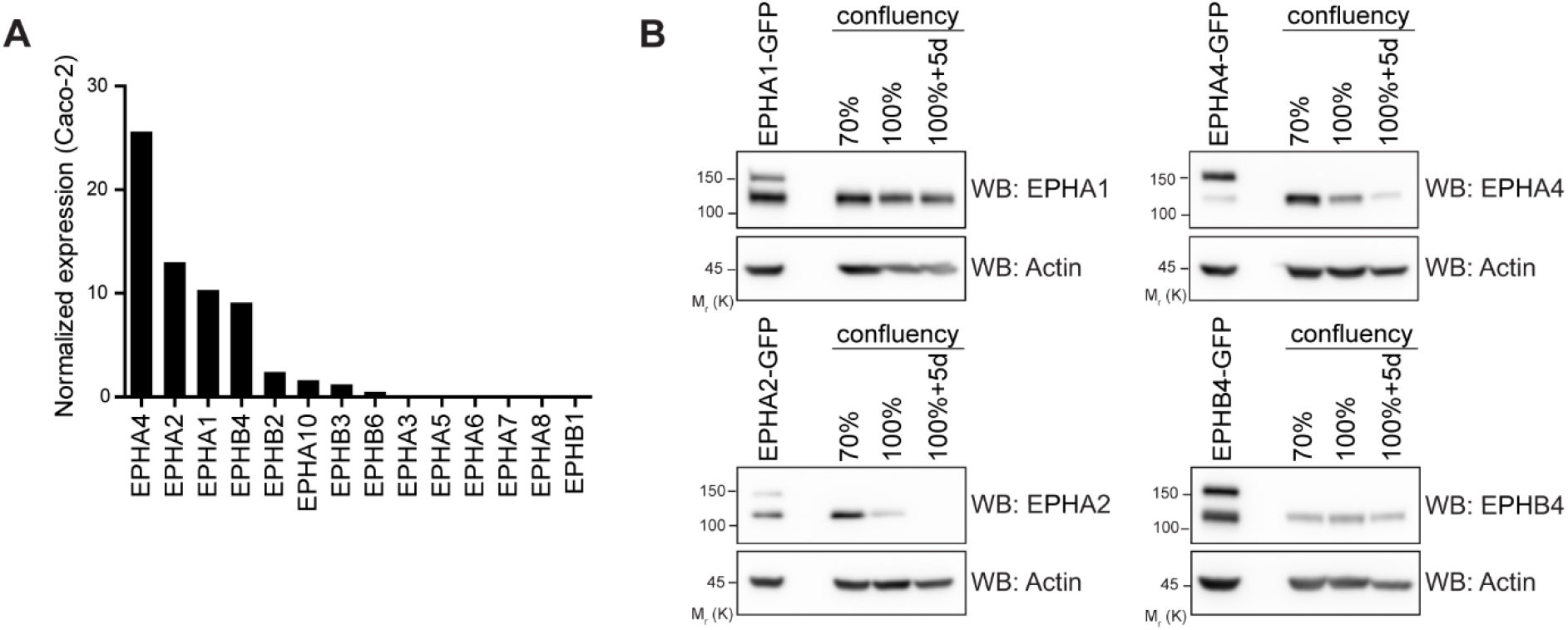
EPHA1 and EPHB4 expression is sustained in polarized Caco-2 cells. (A) mRNA expression levels of all human EPHRs in Caco-2 cells were extracted from the Human Protein Atlas database and displayed in a decreasing order. (B) Western Blot analysis of endogenous EPHA1, EPHA2, EPHA4 and EPHB4 in sub-confluent (70%), confluent (100%) and post-confluent/polarized (100%+5d) Caco-2 cells cultured as monolayers. Overexpressed GFP-tagged EPHRs are utilized as antibody controls. Representative images of 3 experiments are shown.

**Figure S4.**
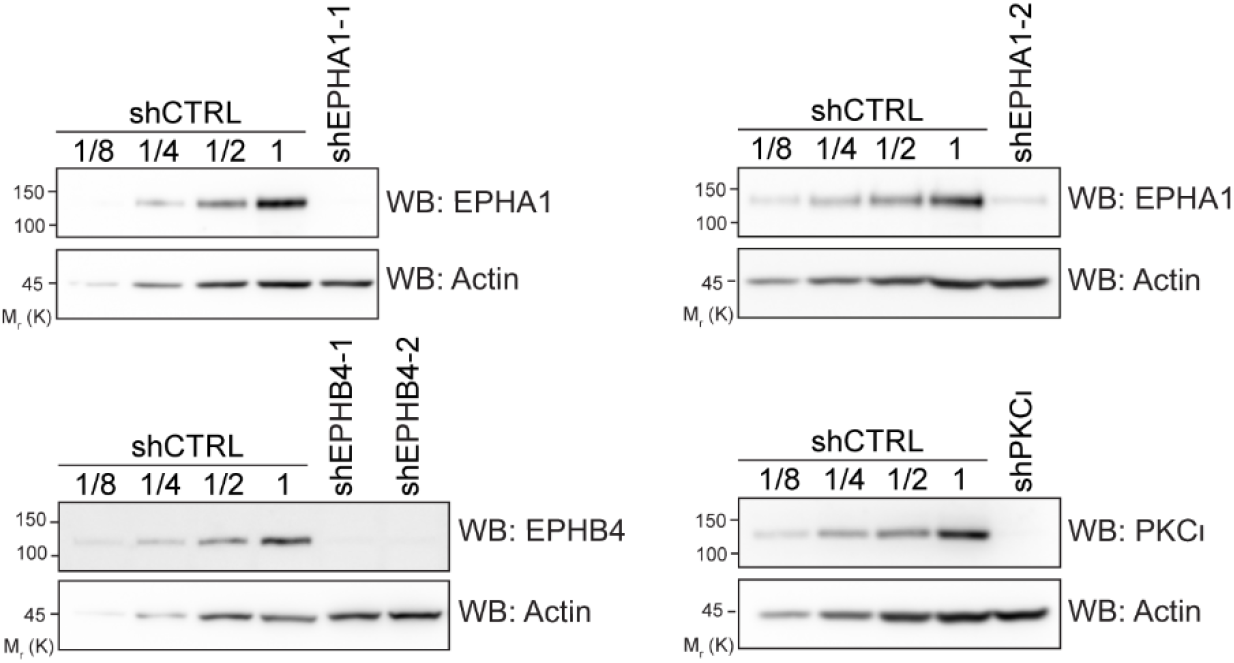
Endogenous PKCι, EPHA1 or EPHB4 depletion. Western blot analysis of endogenous PKCι, EPHA1 or EPHB4 following shRNA depletion for experiments shown in Figure 3. Successive dilutions are indicated to estimate depletion efficiency. Representative images of 3 experiments are shown.

**Figure S5.**
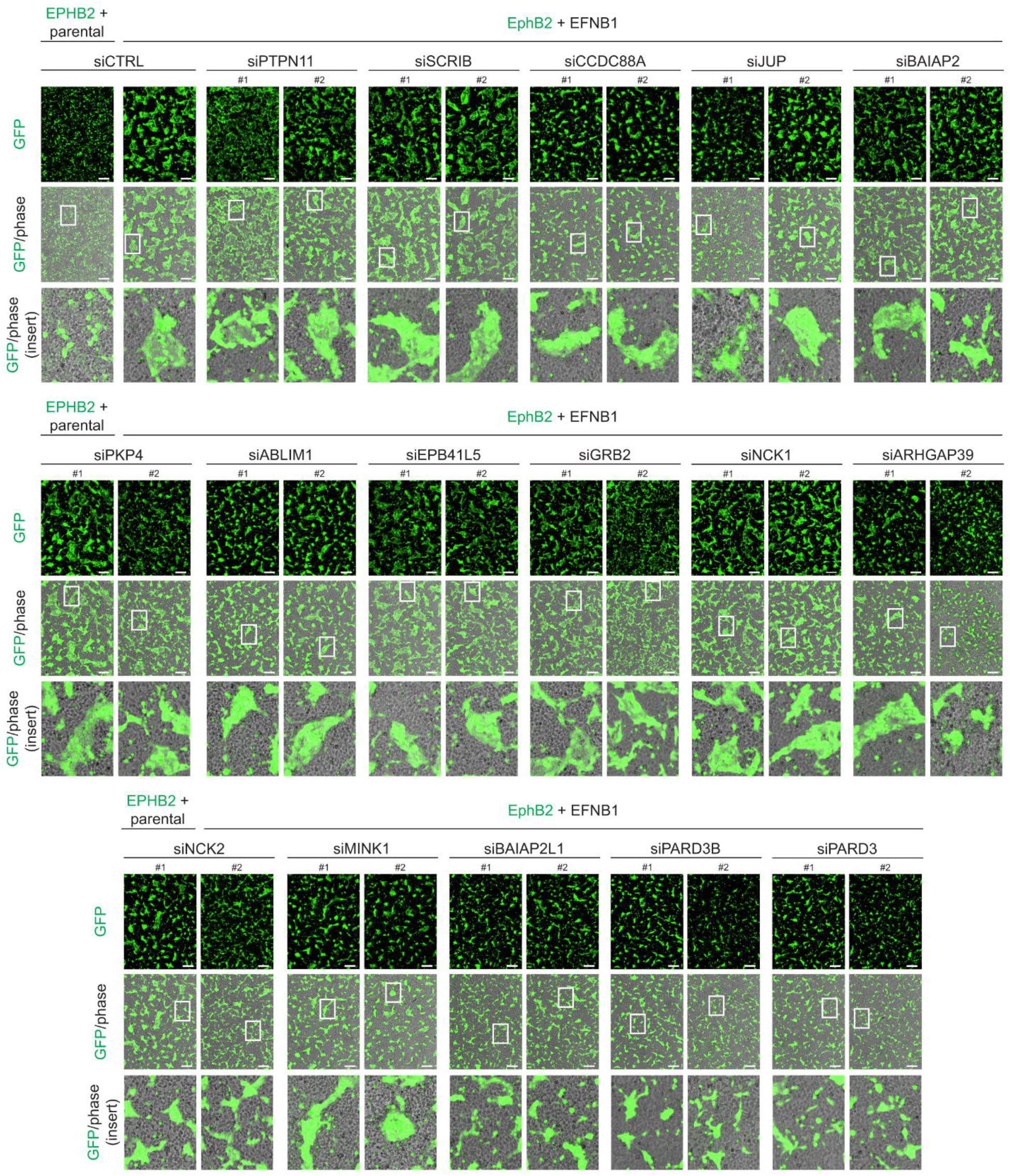
Effect of candidate loss-of-function on EPHR-dependent cell segregation. Representative images (Figure 4C) of cell co-cultures following siRNA depletion of selected candidates in EPHB2 expressing cells and mixing with ephrinB1-expressing cells, visualized by GFP (scale bar: 250 µm).

**Figure S6.**
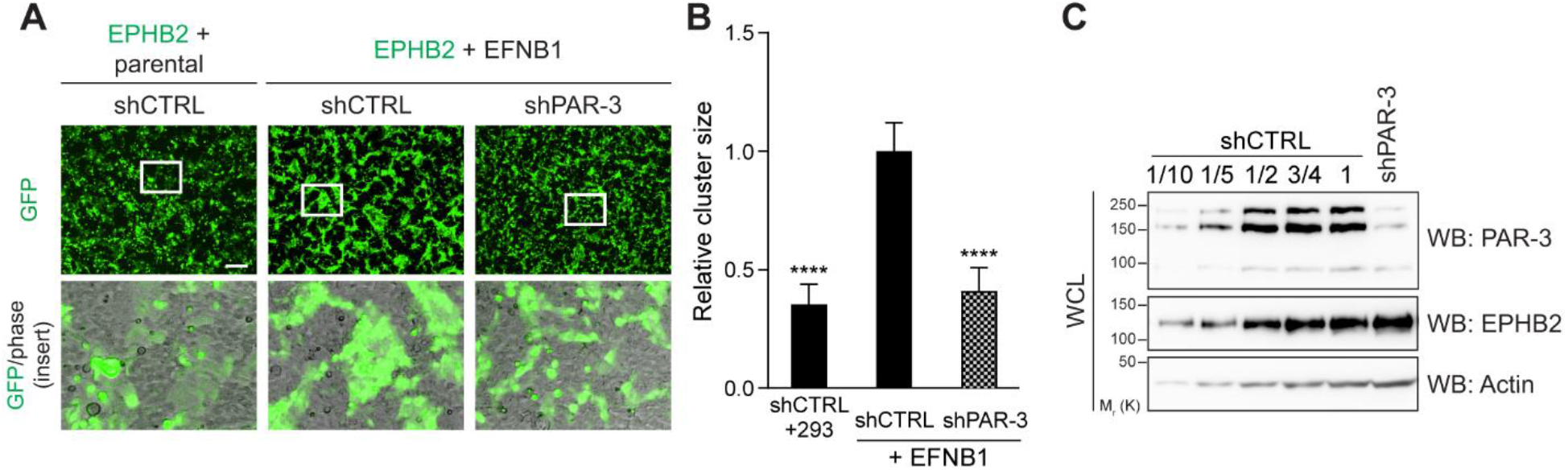
PAR-3 shRNA-mediated depletion blocks EPHR-dependent cell segregation. (A) Representative images of cell co-cultures following shRNA-induced PAR-3 depletion in EPHB2 expressing cells and mixing with EFNB1-expressing cells, as visualized with GFP (scale bar: 250 µm). (B) Quantification of the average size of EPHB2-GFP cell clusters depleted with a shRNAs targeting PAR-3, following mixing with ephrinB1-expressing cells, relative to control (EPHB2 siCTRL cells mixed with ephrinB1 cells). The mean of three experimental replicates is shown. Error bars indicate SD (**** p < 0.0001; unpaired t-test). (C) Western blot analysis of endogenous PAR-3 following shRNA depletion. Successive dilutions are indicated to estimate depletion efficiency. Representative image from 3 experiments is shown.

**Figure S7.**
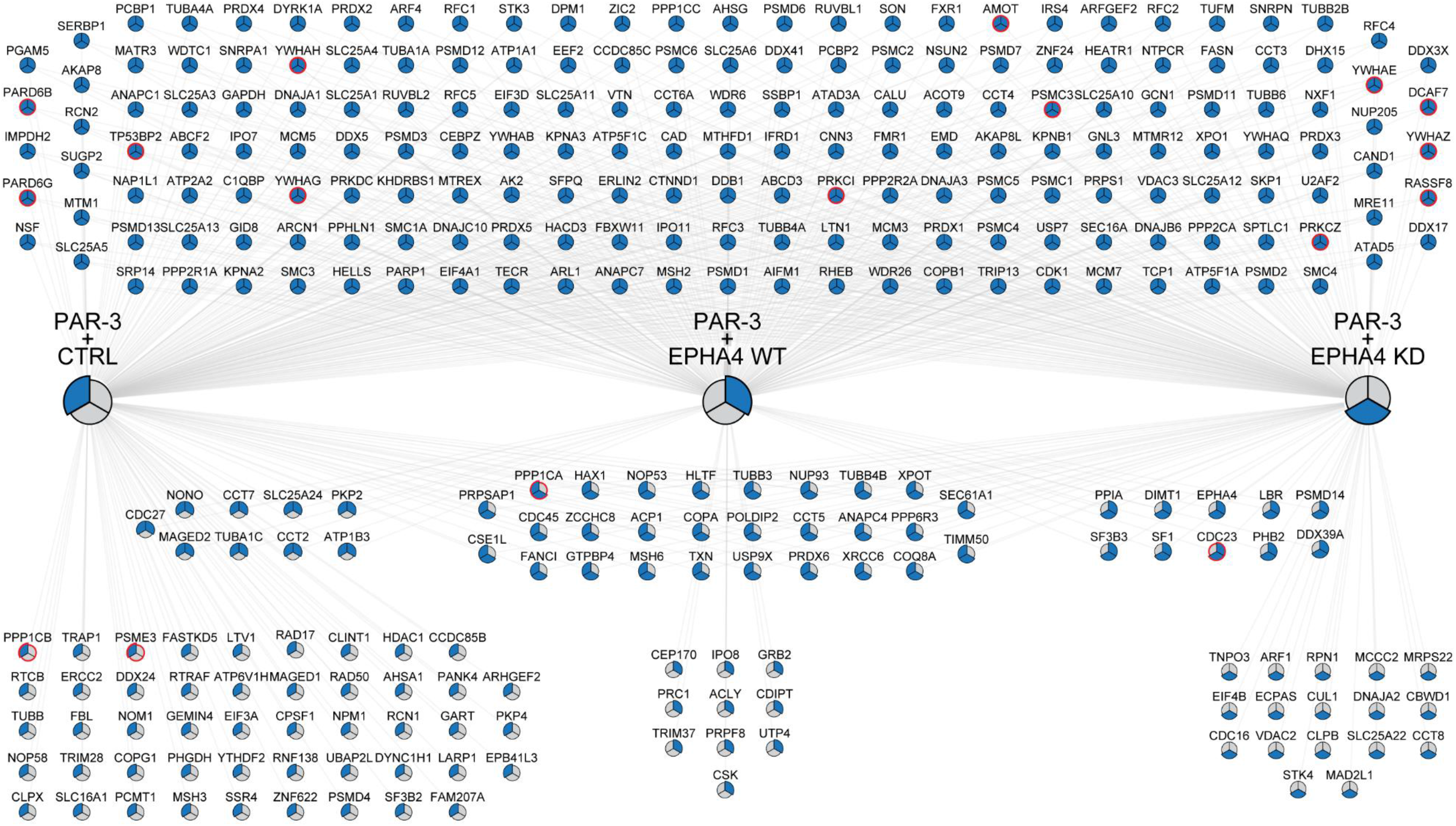
The PAR-3 interactome is modulated by EPHA4. *Bona fide* components of the PAR-3 interactome in the absence/presence of EPHA4 WT or KD, following the removal of background contaminants and non-specific interactions via SAINTexpress using a GFP control (BFDR ≤1%). AP-MS experiments were performed in triplicate. Ribosomal proteins and ribonucleoproteins were further manually removed. Candidates previously reported in the BioGRID database for each EPHR bait are noted with a red edge.

## SUPPLEMENTAL DATASETS

**Table S1:** Contains results of SAINTexpress analyses of EPHA4, -B2, -B3, -B4 BioID experiments (Excel document). Related to Figures 2A, S1 and S2.

**Table S2:** Contains results of SAINTexpress analyses of PAR-3 AP-MS experiments (Excel document). Related to Figures 6A and S7.

**Table S3:** Contains a list of DsiRNA sequences used in cell sorting experiments (Excel document). Related to Figures 4B-C, 7A-C and S5.

**Table S4**: Contains PAR-3 peptide sequences used in *in vitro* kinase array and results of the image quantification (Excel document). Related to Figure 5E.

**Mass spectrometry data availability:** The mass spectrometry proteomics data have been deposited to the ProteomeXchange Consortium (ProteomeXchange ID: PXD021257) via the MassIVE partner repository (http://massive.ucsd.edu), with the dataset identifier MSV000086054 (with the password “polarity” to access files prior to publication). Additional files include a list of BioID and AP-MS experiments, controls, and sample files, the complete SAINTexpress outputs for each dataset as well as a ‘‘README’’ file that describes the experimental procedures.

